# Sculpting conducting nanopore size and shape through *de novo* protein design

**DOI:** 10.1101/2023.12.20.572500

**Authors:** Samuel Berhanu, Sagardip Majumder, Thomas Müntener, James Whitehouse, Carolin Berner, Asim K. Bera, Alex Kang, Binyong Liang, G Nasir Khan, Banumathi Sankaran, Lukas K. Tamm, David J. Brockwell, Sebastian Hiller, Sheena E. Radford, David Baker, Anastassia A. Vorobieva

**Affiliations:** Department of Biochemistry, The University of Washington, Seattle, WA, USA; Institute for Protein Design, University of Washington, Seattle, WA, USA; Biozentrum, University of Basel, Basel, Switzerland; Astbury Centre for Structural Molecular Biology, School of Molecular and Cellular Biology, Faculty of Biological Sciences, University of Leeds, Leeds LS2 9JT; Structural Biology Brussel, Vrije Universiteit Brussel, Brussels, Belgium; VUB-VIB Center for Structural Biology, Brussels, Belgium; Department of Molecular Physiology and Biological Physics and Center for Membrane and Cell Physiology, University of Virginia, Charlottesville, VA 22903, USA; Molecular Biophysics and Integrated Bioimaging Division, Lawrence Berkeley National Laboratory, Berkeley, CA, USA; Howard Hughes Medical Institute, University of Washington, Seattle, WA, USA; VIB Center for AI and Computational Biology, Belgium

## Abstract

Transmembrane β-barrels (TMBs) are widely used for single molecule DNA and RNA sequencing and have considerable potential for a broad range of sensing and sequencing applications. Current engineering approaches for nanopore sensors are limited to naturally occurring channels such as CsgG, which have evolved to carry out functions very different from sensing, and hence provide sub-optimal starting points. In contrast, *de novo* protein design can in principle create an unlimited number of new nanopores with any desired properties. Here we describe a general approach to the design of transmembrane β-barrel pores with different diameter and pore geometry. NMR and crystallographic characterization shows that the designs are stably folded with structures close to the design models. We report the first examples of *de novo* designed TMBs with 10, 12 and 14 stranded β-barrels. The designs have distinct conductances that correlate with their pore diameter, ranging from 110 pS (∼0.5 nm pore diameter) to 430 pS (∼1.1 nm pore diameter), and can be converted into sensitive small-molecule sensors with high signal to noise ratio. The capability to generate on demand β-barrel pores of defined geometry opens up fundamentally new opportunities for custom engineering of sequencing and sensing technologies.

**One sentence summary:** De novo design enables the generation of stable and quite transmembrane beta-barrel nanopores with tailored sizes, shapes and properties.

## Main text

Transmembrane β-barrel (TMBs) nanopores formed by a circularly-closed single β-sheet provide rigid scaffolds for the transport of molecules across cellular (e.g. pore-forming toxins (*1*)) and organelle membranes (outer membranes of bacteria (*2*), mitochondria (*3*) and chloroplast (*4*)). Engineering of naturally occurring nanopores has enabled single-molecule enzymology (*5*), protein fingerprinting (*6*), the detection of small molecules and biomarkers (*7*), and the sequencing of biological and synthetic polymers (*8*). Of particular note is nanopore-based DNA sequencing (*9*), which has enabled widely-accessible large-scale genomics, epigenomics and microbiological analysis (*10*). Despite this success, the development of nanopore sensors for robust analysis of molecules beyond DNA sequencing has so far been challenging. The sensing properties of a nanopore for an analyte of interest can be modulated by introducing mutations into the pore lumen that alter nanopore/analyte interactions (*11*). It remains however challenging to identify a channel suitable for each of the many applications of interest, because there is only a limited set of engineerable naturally occuring nanopores, and these have mostly evolved for very different functions. Going beyond nature, a conducting pore based on a β-hairpin peptide has been designed that transports poly-lysine peptides (*12*). Such self-assembling β-hairpins are however not suitable as a general approach to nanopore design, because it is challenging to control the channel size and to assemble the pore in lipid membranes. Monomeric 8-stranded TMBs have been designed that stably assemble in detergent and in lipid vesicles, but they are too small to contain a central conducting channel (*13*).

Encouraged by the success designing these narrow TMBs, we reasoned that *de novo* protein design should provide a general approach to creating robust β-barrel nanopore scaffolds for a next generation of nanopore sensors. A key challenge in designing such structures is that the polar-hydrophobic pattern characteristic of globular protein folds must be inverted: the exterior must be largely nonpolar for membrane insertion, and the interior must be largely polar to support a solvated conducting channel. Furthermore, unlike globular proteins, the structure of TMBs must be specified in the vast majority by short-range interactions between residues located on adjacent strands since there is no close-packed core. Finally, the amphipathic β-strands are highly aggregation prone prior to β-barrel assembly, and hence the design must strongly favor intra-chain rather than inter-chain interactions during folding. We set out to develop general methods to overcome these challenges and design stable monomeric channels with tunable pore shapes, sizes, and single-channel conductance.

### Computational design

We sought to build from scratch TMB backbones accommodating water-accessible pores starting from the principles elucidated during the design of 8-stranded TMBs lacking pores (*13*). To modulate the size of the pore, we increased the number of β-strands (10, 12 and 14 strands) while keeping the transmembrane span and the connectivity between β-strands (the shear number (*14*, *15*)) constant. This resulted in an increase of the average β-barrel diameter from 16.4 Å for the previously designed 8 strand β-barrels (*13*) to 19.4 Å (10 strands), 22.8 Å (12 strands) and 26.4 Å (14 strands) (Figure 1A, Figure S1). By comparison to 8-stranded TMBs, the diameters of the larger β-barrels do not allow long-range sidechain contacts across the pores and the structural properties of the pores (β-strand pairing, β-barrel shape) must be locally encoded. Naturally-occurring TMBs typically feature long, disordered, loops on one side of the barrel (*16*), which results in noisy electrophysiology recording and challenging data interpretation when the pores are used for sensing applications (*17*, *18*). To design quiet pores and reduce the noise, we connected the β-strands on both sides of the barrels with 2- and 3-residues (Figure S2) β-hairpins, the shortest loops we have previously found to support TMB folding (*13*). The first-generation backbones corresponding to these designs were assembled with the Rosetta BlueprintBDR (*19*) application and had similar cylindrical shapes. Such cylindrical β-sheet configurations are strained (*20*, *21*) due to repulsion between side-chains packing the barrel lumen (Figure 1C). Glycine kinks (glycine residues in extended positive-phi conformation (*15*)) were introduced into the blueprint to relieve the strain and to bend the β-strands to form corners in the β-barrel cross-section. We generated four blueprints with the same topology but different glycine kink distributions to design 12-stranded β-barrel backbones with square, triangle, rectangle or oval-shaped cross-sections (Figure S1). A single glycine kink was used in corners of an angle of >= 90° and several adjacent and/or stacked kinks were placed to form corners of < 90° (Figure 1B). Sequence-agnostic TMB backbones incorporating these constraints were assembled *in silico* and had the shapes expected based on the placement of the glycine kinks (Figure 1D).

**Fig. 1:**
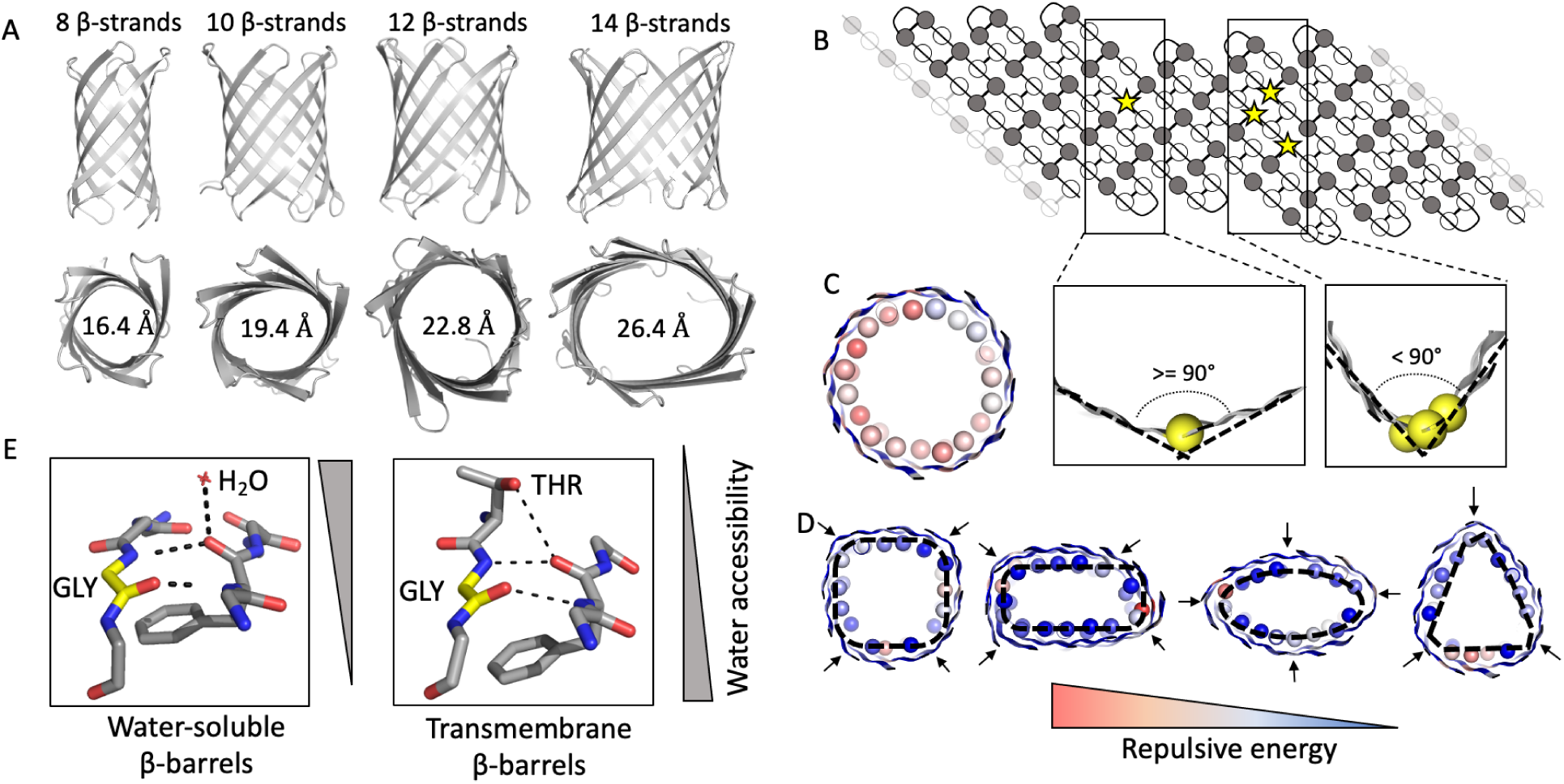
Sculpting β-barrel geometry. A. Pore diameter can be controlled through the number of β-strands in the β-barrel blueprint. B. β-barrel 2D interaction map. Strong bends in the β-strands (< 90° bend, right) is achieved by stacking several glycine kink residues (yellow spheres) along the β-barrel axis, as opposed to placing one kink (>90° bend, left). C-D. Cross-sections of explicitly assembled β-barrel backbones without (cylinder, C) and with (D) glycine kinks. The Cβ atoms of the residues facing the pore are shown as sphere’s and colored based on their respective repulsion energy. Glycine kinks positions are shown with arrows; placement at the corners of the embedded rectangular, oval and triangular shapes (dashed lines in D) generates the desired backbone geometries. E. Polar threonine residues are tolerated on the membrane-exposed surface of TMBs (right) as they can form a hydrogen bond to the backbone, mimicking the interactions with water molecules observed in similarly curved areas of water-exposed β-strands (left).

A challenge for TMB design is to balance the optimization of the folded β-barrel state in the membrane with delayed folding in water to reduce misfolding and aggregation that would prevent successful integration into a membrane bilayer (*13*, *22*, *23*). For the 8-stranded TMBs, this was achieved by incorporating local secondary-structure frustration (*24*) to reduce premature formation of aggregation-prone β-strands prior to full barrel assembly: hydrophobic amino acids were designed into the water-accessible pore to disrupt the hydrophobic-polar amino acid alternation pattern characteristic of amphipathic β-sheets. To test whether such balancing is necessary for larger β-barrel designs that need to have water-accessible (and hence more polar) channels, we first designed “optimal” 10 and 12-strand TMBs with only polar and charged amino acids facing the pore. All 16 such designs failed to express in *E. coli*, as was previously observed for similarly optimal 8-stranded TMB designs. We therefore set out to design larger TMB nanopores incorporating local secondary structure frustration. In the water-accessible pore, networks of polar residues were designed around the canonical TMB folding motif Tyr-Gly-Asp/Glu (*13*, *25*, *26*) to optimize strong local β-register defining interactions while alternating with patches of hydrophobic and small, disorder promoting, residues (Gly, Ala, Ser). To compensate for the more hydrophilic pores in the larger TMBs, we further reduced the β-sheet propensity by replacing a small number of β-branched residues with Ser and Thr amino acids on the lipid-exposed surface. Although it is perhaps counterintuitive to expose hydroxyl groups to the lipid environment, model building indicated that Ser and Thr could form a hydrogen bond to the β-strand backbone when placed in close proximity with a glycine kink, effectively mimicking the backbone-water hydrogen bonds observed in strongly bent β-strands of water-soluble β-barrels (Figure 1E).

During combinatorial design of sequences for β-barrels of different size, we found that the frequency of incorporation of each amino acid type strongly depended on the curvature of the β-sheet. For each of the generated blueprints, we adjusted the Rosetta solvation and reference energies (*27*) to achieve the desired balance of frustrated and energetically-favorable contacts (Figure S3). Following several iterations of combinatorial sequence design and structure relaxation, designs were selected based on hydrogen bond network descriptors, secondary structure (*28*) and aggregation propensities (*29*) (Figure S4). We previously found that AlphaFold2 (*30*) could accurately predict the structures of designed TMBs even in the absence of evolution information (from a single sequence input and without a multiple sequence alignment (*31*)) when weak 3D contacts contained in the sequence were amplified by 48 rounds of molecular model recycling through the prediction network, and that the confidence assigned to the model (plDDT) was a good discriminator of the sequences with higher probability of experimentally folding (*32*). Therefore, we selected 4-10 designs per blueprint for which AlphaFold2 predicted high-confidence structures closely matching the design models (Figure S5).

### Experimental characterization of TMB folding

We first tested two sets of TMBs with 10 (four designs) or 12 β-strands with a square cross-section (nine designs). Genes were synthesized and the proteins were expressed as inclusion bodies in *E. coli* to avoid the complexity of targeting the outer membrane (*33*) (Figure 2A). Unlike the 16 “optimal” designs which all failed to express, most sequences incorporating secondary structure frustration were expressed at high-levels (12/13, Figure S6). Since most naturally-occurring TMBs can fold *in vitro* (*34*), the purified designs were solubilized in guanidine hydrochloride and refolded by slow dilution into a buffer containing either detergent (fos-choline 12 (DPC) at a concentration double the critical micellar concentration (CMC)) or synthetic lipid vesicles (Material & Methods). As previously observed for the 8-stranded TMB designs, the standard band-shift assay on cold SDS-PAGE used to assess folding of natural TMBs (*35*) was not informative to identify properly folded synthetic TMBs (Figure S7). Instead, the designs were characterized by size exclusion chromatography (SEC), far UV circular dichroism (CD) in the presence of DPC detergent, and tryptophan fluorescence in DUPC (C_11:0_PC) large unilamellar vesicles (LUVs). One 10-strand design (TMB10_163) and one 12-strand design (TMB12_3) with predominantly monomeric SEC profiles (Figure 2A), thermostable CD spectra characteristic of β-sheet (Figure 2B,C) and clear shift of tryptophan fluorescence maximum from ∼350 nm (unfolded proteins in 8 M urea or in the absence of lipid) to ∼330 nm (folded in LUVs) (Figure S8, S9) were selected for further characterization by urea titration. Both designs showed sharp and reversible folding/unfolding transitions in the presence of DUPC LUVs (Figure 2D) (mid-point urea concentrations for folding (Cm^F^): 4.5 ± 0.2 M and 5.5 ± 0.2 M, respectively). The equilibrium unfolding curves were fitted to a two-states transition, with the calculated unfolding free energies (ΔG^0^) of -35.6 ± 2.7 and -63.1 ± 8.0 kJ/mol (for TMB10_165 and TMB12_3, respectively) in the range of natural (ΔG^0^ -10 to -140 kJ/mol (*36*–*39*)) and previously designed 8-stranded TMBs (-38 and -56 kJ/mol (*13*)). To confirm that the designs folded by integration into the bilayer rather than partial folding on its surface, the kinetics of folding were recorded in DUPC (C_11:0_PC) membranes, as well as in thicker DMPC (C_14:0_PC) membranes. Dramatically decreased folding rates were observed with DMPC compared with DUPC LUVs (Figure S10), consistent with integral membrane folding. Encouraged by these results, we assessed the nanopore activity of these two designs, based on their capacity to integrate spontaneously into planar dipalmitoylphosphatidylcholine (DPhPC) membranes after dilution out of DPC micelles, and to conduct Na^+^ and Cl^-^ ions. The 12-strand TMB12_3 inserted successfully into the membrane, producing distinct jumps of current of reproducible intensities (Figure S11), and had stable nanopore conductance. While the design TMB10_163 did not have detectable nanopore activity, one variant (TMB10_165) featuring seven mutations on the lipid-exposed surface (V72I, V102T, V114I, A124L, I126V, I138V and I144V, designed using Rosetta (*40*)) inserted successfully into DPhPC membranes and was able to conduct ions (Figure S11). TMB10_165 had increased stability by comparison to TMB10_163, as shown by higher stability to protease digestion and more dispersed NMR ^1^H-^15^N HSQC chemical shift in DPC micelles (Figure S12). Unlike some native pores which can fall off from the membrane stochastically, these two pores remain stably inserted over long periods of time with the longest recording acquired being 2 hours for the TMB12_3 design. Recording of the current-to-voltage response showed an incremental increase in observed conductance under positive and negative changes of voltage, indicative of stable transmembrane channels (I/V curves in Figure S11). Our results on TMB10_163, TMB10_165, TMB12_3 and other TMB12 designs with less, to no, detectable nanopore activity (Figure S14) suggest that membrane integration and nanopore conductance require stable TMB folding *in vitro*.

**Fig. 2:**
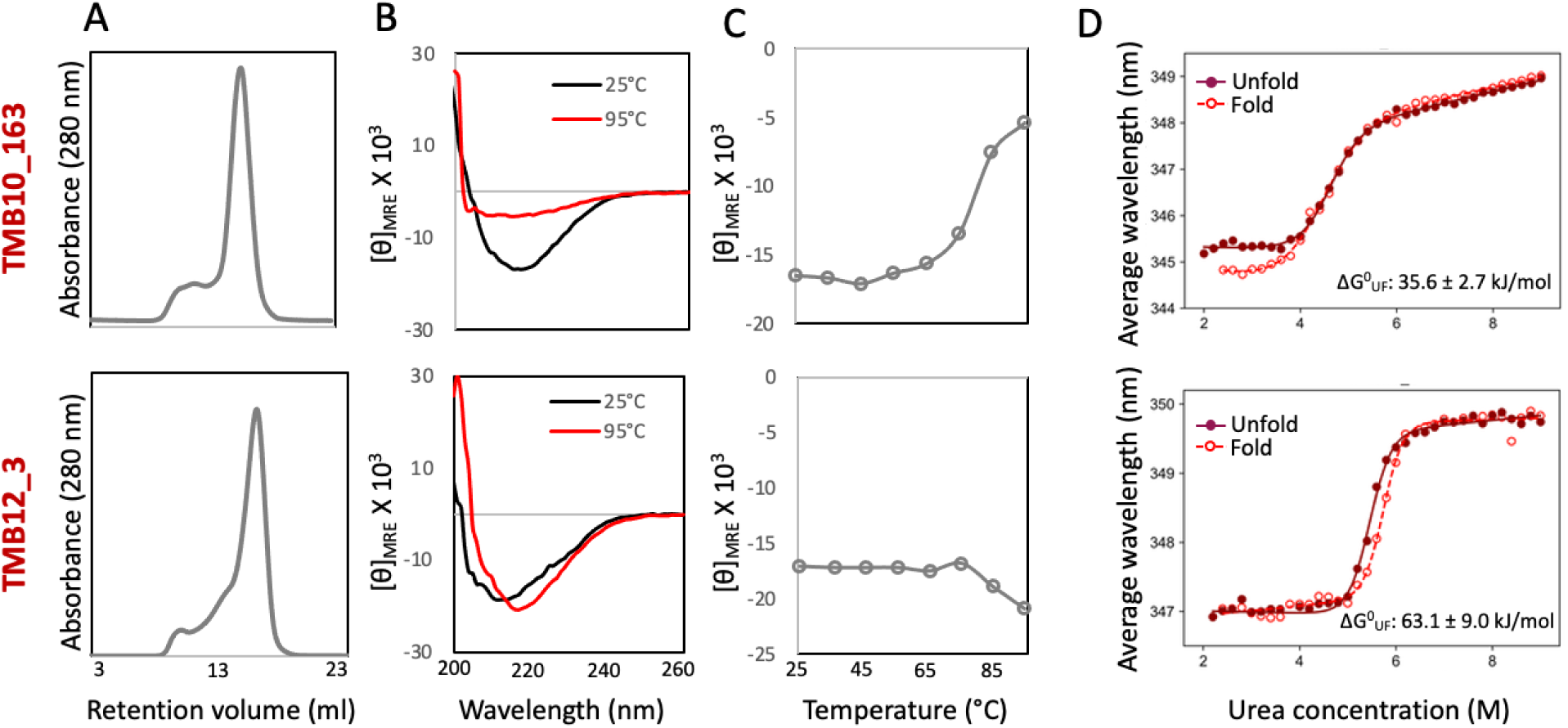
Biophysical characterization of designed nanopores. Top row: 10-stranded design (TMB10_163); Bottom row: 12-stranded design with a square cross-section (TMB12_3). Both designs elute as one major species with retention time consistent with a monomeric protein in complex with DPC detergent (A), show distinct negative maxima in far UV CD spectra at 215 nm (B) that remain stable up to >70°C (D), and cooperative and reversible folding/unfolding transitions in DUPC LUVs (obtained at 25°C) (D),

We next sought to solve the structures of the designs to assess the accuracy of the computational design methods. While the design TMB10_165 did not form crystals in the conditions screened, TMB10_163 formed crystals which diffracted to 2.5 Å resolution (Table S1). The seven surface-exposed mutations between TMB10_165 and TMB10_163 are shown in Figure 3A. The four copies of the TMB10_163 in the asymmetric units had a structure very similar to the original Rosetta design, with an average RMSD of 1.4 Å over all backbone heavy atoms (Figure 3B) and featured the expected β-strand connectivity (shear number of 12). Although TMB10_163 nanopore activity was not observed, analysis of its structure using PoreWalker (*41*) and MOLE 2.5 (*42*) indicated the presence of a water-accessible cylindrical pore with an average diameter ranging from 4.2 to 5.3 Å in the four subunits (Figure 3C, Figure S15), matching the diameter of the pore calculated from TMB10_163 design model (4.6 Å). Most of the side-chains lining the pore had similar rotameric states in the crystal structure and the design model, with remarkable similarity at the level of the designed Tyr-Gly-Asp/Glu folding motifs (Figure 3D). We further determined the structure of TMB12_3 by NMR spectroscopy. Optimization of the *in vitro* folding conditions showed that the protein was structured in aqueous solution in LDAO detergent micelles, as indicated by well-dispersed amide and side chain methyl spectra (Figure 3E, Figure S16). Secondary chemical shifts indicated the presence of twelve β-strands arranged into polypeptide segments expected from the design (Figure S17). Amide and side chain methyl NOEs spanned a dense network of experimental connectivities that reached around the barrel circumference and thus confirmed the correct arrangement of the strands into the predicted barrel structure (Figure 3F). The TMB12_3 features the designed β-strands connectivity (shear number of 14) with the barrel closed by the canonical antiparallel β1-β12 seam (Figure 3G, Figure S18, Table S2). These experimental structures demonstrate that our computational design method can design TMB nanopores with precisely controlled shear, channel width and shape.

**Fig. 3:**
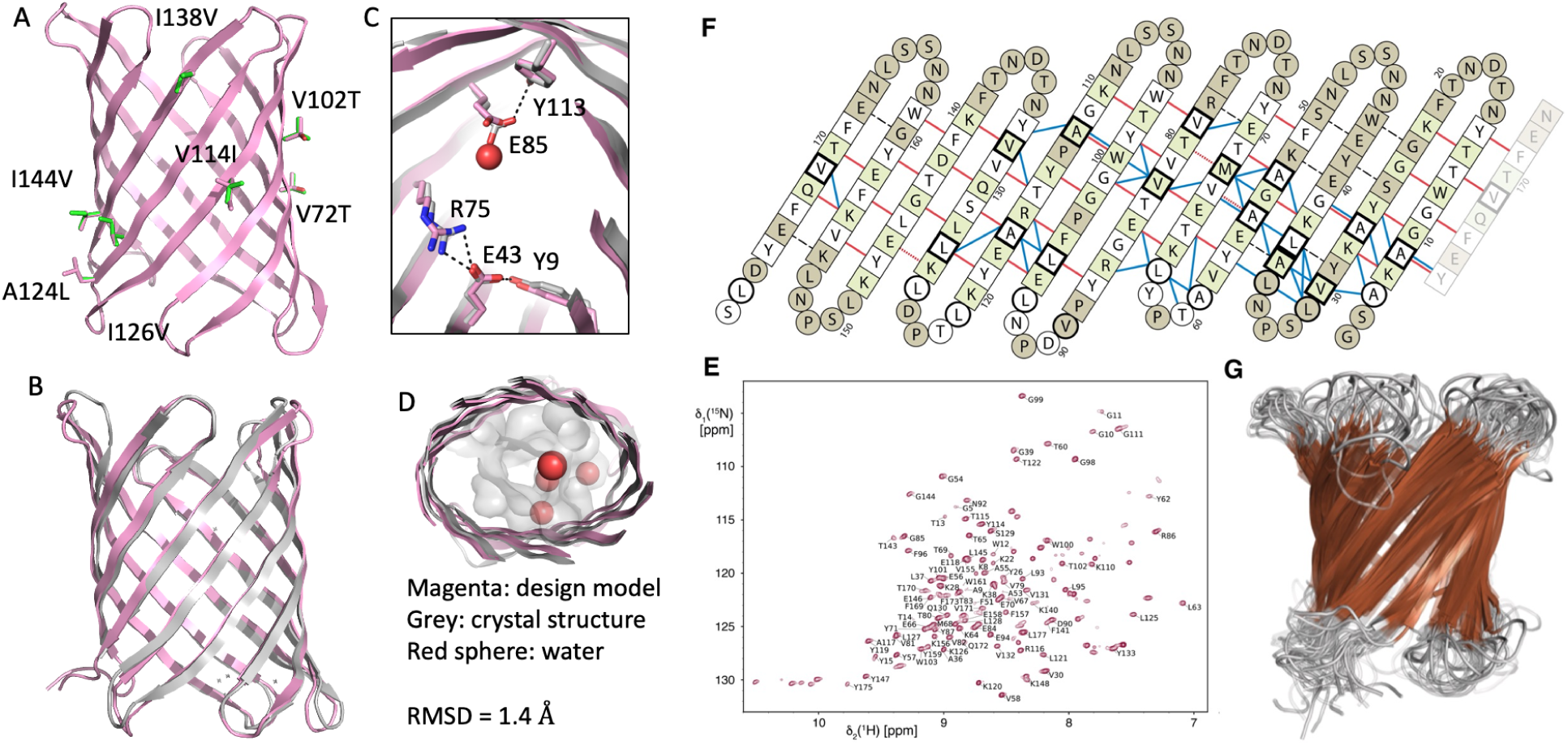
Experimentally determined nanopore structures closely align with the computational design models. A. Seven surface mutations differentiate TMB10_163 from TMB10_165. B-D. Crystal structure of TMB10_163 B. backbone superposition C. superposition of side-chains involved in key folding motifs in the lumen D. cross-sections superposition with the water-accessible pore shown as a gray surface. Water-molecules crystallized in the pore are shown as red spheres. E-G. TMB2_13 structure in LDAO micelles. E. 2D [^15^N,^1^H]-TROSY NMR spectrum of [U-^2^H,^15^N]-TMB12_3 with sequence-specific resonance assignments. F. Long-range NMR NOE contacts mapped to the expected TMB12_3 hydrogen bonds (dashed black lines). Residues with amide assignment are shown in white and green, unassigned residues are shown in ash gray. Residues with β-sheet secondary structure are shown as squares, all others as circles. Bold outlines indicate available methyl assignments. NOE contacts are shown as red lines (long-range amide-amide, dashes indicating diagonal overlap) and blue lines (contacts involving side chain methyl groups). G. Ensemble of the 20 lowest energy solution NMR structures (β-sheets shown in brown).

### Electrophysiology

Encouraged by the success in designing 10- and 12-stranded β-barrels, we set out to design TMBs with different numbers of β-strands and different shapes. We designed 12-stranded β-barrels with a triangular cross-section (eight designs), an oval cross-section (seven designs), or a rectangular cross-section (nine designs), as well as 14 β-stranded β-barrels (nine designs), incorporating the design features described above for the 10- and 12-stranded TMBs. The designs were obtained as synthetic genes and the proteins were again expressed in inclusion bodies. A relatively lower fraction of 12-stranded TMB designs with a rectangular (4/9 designs) and oval (4/7 designs) cross-section showed a prominent expression band SDS-PAGE gel in comparison to the square-shaped designs (8/9). This could be the result of less homogeneous distribution of β-sheet destabilizing amino acids (which are easier to introduce in bent than in flat β-sheet regions) in these designs, as suggested by a higher density of strong β-sheet islands co-localizing with predicted early folding regions (*43*) (Figure S19). The difficulty of *de novo* β-barrel design thus depends not only on the size of the TMB pore but also on the shape encoded into the blueprint.

Because of the small number of designs pre-selected *in silico* thanks to the new AlphaFold2 filter, we proceeded to screening designs for nanopore activity directly after validating their folding into monodispersed β-sheet structures in DPC micelles (Figure S20). We evaluated the designs for their capacity to insert into planar membranes from dilute detergent solution and form conducting pores (Figure 4). We obtained both 12 (three triangular-shaped, three oval-shaped and two rectangle-shaped) and 14 stranded (two) TMBs that exhibited consistent and stable conductances at positive and negative voltage (Fig 4, 3rd and 5th columns), with multiple sequential insertions corresponding to current jumps of small integral multiples of the base pore conductance (Figure 4, 4th column). Based on the intensities of the current jumps, we estimated the conductances of single-channel events, which increased with pore size, as expected: the 10 stranded TMB design described above had a conductance of 108 ± 1.4 pS, which based on the cylindrical pore access resistance model (*44*) corresponds to a nanopore diameter of approximately 3.5 Å. The 12-stranded designs had similar conductances to each other (210-230 pS) despite their different shapes, consistent with a cylindrical nanopore of around 5 Å. The 14-stranded design had a conductance of 427 ± 2.7 pS consistent with a calculated pore diameter of 7 Å. The predicted diameters are close to the average expected diameters of 4.6 ± 0.7 Å, 9.4 ± 0.8 Å and 10.6 ± 1.4 Å (calculated along the pore of TMB10_165, TMB12_3 and TMB14_8 design models, respectively, using MOLE 2.5 (*42*) (Figure S15)). In comparison to naturally-occurring pores used for sensing, such as OmpG which undergoes both transient and complete occlusion events by its solvent-exposed loops over a timescale of 100 ms (*18*, *45*), our TMB designs show remarkably quiet conductances, with no occlusion events detected over 10 sec measurements (Figure S11). Varying the pore shape of the pore while keeping the size constant (Figure 4, first column) did not have a large effect on monovalent ion conductance, likely because of the relatively large pore constrictions and the flexibility of the long polar side-chains lining the channels (Figure S21). Nevertheless, we anticipate that modulation of the nanopore shape and chemical lining should allow control over the permeability of the pores to larger and more complex solutes in the future.

**Fig. 4:**
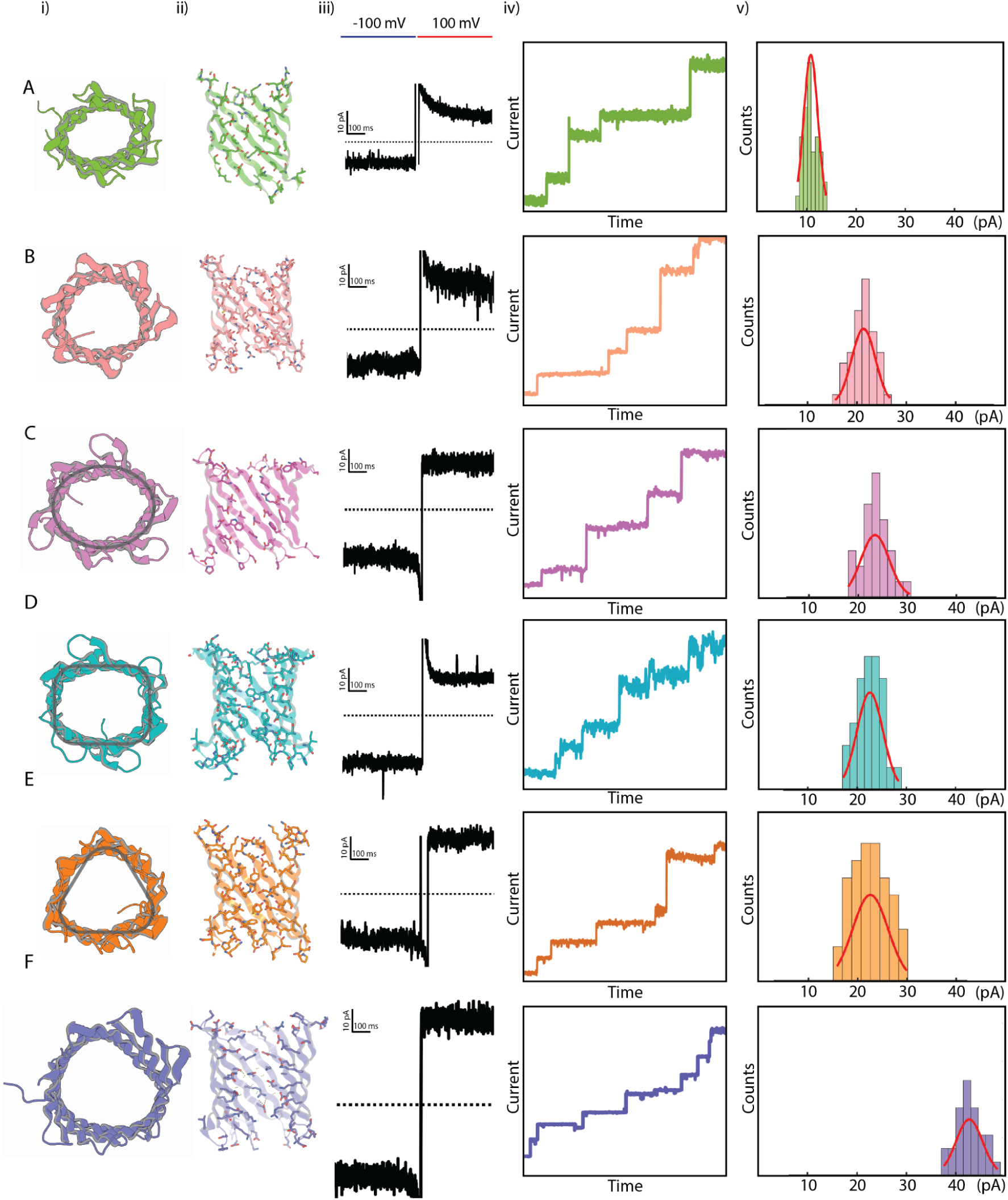
Conductance of designed nanopores. A. TMB10_165, B. TMB12_3, C. TMB12_oval_4, D. TMB12_rect_8, E. TMB12_tri_12, F. TMB14_8. i) Top view cartoon representation. ii) Vertical cross sections of the pore. iii) single channel conductance (smallest observed conductance jump). iv) sequential insertions of designed pore in planar lipid bilayer membrane from detergent solubilised sample at low concentrations. v) histogram of smallest measured current jumps for each design up to 50 pA. The applied voltage across the bilayer was 100mV and experiments were performed in a buffer containing 500 mM NaCl. A gaussian fit was carried out for the single channel current histograms for each design. For TMB10_165, 38 independent single channel jumps were identified from 3 recordings to plot the histogram shown. Similarly, 44 single channel insertions were identified for TMB12_3 (4 recordings), 29 insertions for TMB12_oval_4 (3 recordings), 30 insertions for TMB12_rect_8 (3 recordings), 45 insertions for TMB12_tri_12 (5 recordings) and 32 insertions for TMB14_8 (3 recordings) to plot the above depicted histograms.

## Conclusion

Our results demonstrate that it is now possible to systematically design transmembrane β-barrels with conducting pores spanning a range of sizes and shapes. Despite the inversion of the hydrophobic-polar core compared to globular proteins, and the almost entirely local nature of the sidechain interactions, our approach enables TMB design with atomic level precision, as highlighted by the close agreement between the experimentally determined crystal and NMR structures and the corresponding design models. Whereas the shapes of globular proteins are largely determined by the packing of hydrophobic residues in a central core, the TMB shapes can be specified by strategic placement of glycine residues at which bending takes place to reduce strain. As previously observed for 8-stranded TMBs, a delicate balance between the optimization of tertiary structure energy and negative design (introduction of locally frustrated residues) to disfavor premature β-strand formation before membrane insertion was critical for the expression of the larger TMB nanopores in *E coli* inclusion bodies. In comparison with previously designed oligomeric protein nanopores - built from self-assembling ɑ-helical (*46*, *47*) or β-hairpin peptides (*12*) - the nanopores presented here have the advantage of being built from a single chain which enables controlled assembly of monodisperse nanopores without alternative oligomeric states and much greater control over the shape and specific surface properties of the transmembrane channel. While nanopore designs from oligomeric β-hairpins require lipid nanoparticles for solubility and assemble under electric current at lipid-lipid interfaces (*12*), the monomeric TMB designs fold efficiently into detergent micelles and lipid vesicles. The stability of the monomeric designs enables their spontaneous insertion into planar lipid membranes following dilution out of detergent micelles, opening the door to the use of synthetic transmembrane nanopores in commercial flowcells developed for single-molecule sensing and sequencing.

Monomeric integral TMBs such as OmpG have been turned into sensors by incorporating analyte-recognition motifs (*48*, *49*) or biotin-bound (*50*, *51*) antibodies in the solvent-exposed loops (*7*). However, the long disordered loops of naturally-occurring TMBs result in noisy reading, and the engineering of quiet pores with shorter or mutated loops has been a long-standing problem (*17*, *18*, *45*, *52*). The TMBs described here were designed with the shortest loops compatible with folding, and allow up to 2 hours of quiet recording which would enable the detection and quantification of molecules at low concentration. As illustrated in the accompanying manuscript, the stability and robustness of the designed nanopore structures and conductances enables ready conversion into ligand gated channels with considerably lower noise and more comprehensible signal analysis than previously engineered channels.

Further advances in understanding the relation between the chemical properties of a nanopore and the directed detection of an analyte in the pore lumen (*11*, *53*, *54*), coupled with our new ability to design custom pores, opens up exciting new directions for sequencing and sensing since the pore size and the side chains lining the pore can be designed specifically for the desired application. Unlike native pores, which are finite in number, there is no limit on the number of distinct designed pores that can be generated. *De novo* design now enables pore geometry and chemistry to be custom built to be optimal for applications ranging from detection and selective transport of a wide range of molecules of interest to biopolymer sequencing.

## Acknowledgments

We thank Jens Gundlach and Andrew Laszlo from the Department of Physics (University of Washington, Seattle) for helpful suggestions regarding conductance measurements, Ines Hertel for protein production, Bob Schiffrin (fitting of CD denaturation data) and many more colleagues at the VUB-VIB Center for Structural Biology, the Institute for Protein Design and the University of Leeds for helpful discussions.

## Author contributions

AAV and DB designed the research. SB, SM and AAV designed nanopore proteins. AAV wrote the design and analysis scripts. SB and SM expressed, purified proteins and ran SEC and CD in detergent. JW performed CD, equilibrium folding/unfolding and folding kinetics in LUVs with help from GNK and supervised by SER and DJB. TM performed initial ^1^H-^15^N NMR screening of designs, optimization refolding conditions and solved the NMR structure of TMB12_3, supervised by SH. AB crystallized and solved the X-ray structure of TMB10_163 with help of AK and BS. BL collected initial NMR data on TMB10 designs, supervised by LKT. CB developed the electrophysiology characterization method and collected initial data. SM collected electrophysiology data on nanopore designs. AAV and DB wrote the first manuscript draft with support from SH, TM and SM. All authors provided input on the final manuscript.

## Data and materials availability

The Rosetta software suite is available free of charge to academic users and can be downloaded from https://www.rosettacommons.org. The scripts and the designed protein models are available from GitHub (https://github.com/vorobieva/demo_TMB_design) and will be archived in Zenodo. Analysis scripts for processing ion conductance data as presented in this manuscript are also available on Github (https://github.com/sagardipm/denovoPores). The crystal structure of the design TMB10_163 and the NMR structure of TMB12_3 have been deposited in the Protein Data Bank (PDB) (XX, 8UZL). Plasmids of the constructs are available upon request to the corresponding authors.

## Supplementary materials

### Material & Methods

#### General purification of all designs

All designs were purified from *E. coli* following a similar protocol as described previously in *Vorobieva, Anastassia A., et al. "De novo design of transmembrane β barrels." Science 371.6531 (2021): eabc8182*. Custom genes in a pET29b vector containing the kanamycin resistance gene were ordered from IDT and chemically transformed into BL21(DE3)-Star cells. All proteins were purified from inclusion body fractions following complete denaturation in 6M GuCl (guanidine hydrochloride) buffer. Briefly, inclusion pellets were washed several times with buffers containing 1% w/v of Triton X-100 and Brij-35 alternatively. A typically washing step involved resuspension of the insoluble pellet in the appropriate buffer, brief sonication and subsequent incubation for one hour at room temperature or overnight at 4°C. After solubilisation of the pellet in GuCl, the protein was diluted to 80-100 µM and refolded in a buffer containing 25mM Tris-Cl at pH 8.0, 150 mM NaCl and 0.1% w/v DPC (dodecyl-phosphatidylcholine) either using a dropwise dilution method or by spontaneous dilution to achieve a final GuCl concentration of 0.3 M. The diluted buffer was concentrated after overnight incubation with shaking at 4°C and run on a S200 Cytiva Superdex 200 column. Fractions at expected elution volume were concentrated with a 10kDa cutoff filter and used for subsequent analysis.

#### Conductance measurement in planar lipid bilayers

All ion-conductance measurements were carried out using the Nanion Orbit 16TC instrument (https://www.nanion.de/products/orbit-16-tc/) on MECA chips. Lipid stock solutions were freshly made in dodecane at a final concentration of 5 mg/mL. DPhPC (di-phytanoyl-phosphatidylcholine) lipids were used for all experiments. Designed proteins were diluted in a buffer containing 0.05 % DPC (∼ 1 CMC), 25 mM Tris-Cl pH 8.0 and 150 mM NaCl to a final concentration of ∼100 nM. Subsequently, 0.5 µL or less of this stock was added to the cis chamber of the chip containing 200 µL of buffer while simultaneously making lipid bilayers using the in-built rotating stir-bar setup. All measurements were carried out at 25*C. Spontaneous insertions were recorded over multiple rounds of bilayer formation. All chips were washed with multiple rounds of ethanol and water and completely dried before testing subsequent designs. A 500 mM NaCl buffer was used on both sides of the membrane for all current recordings. Raw signals were recorded at a sampling frequency of 5 kHz. Only current recordings from bilayers whose capacitances were in the range 15-25 pF were used for subsequent analysis. The raw signals at 5 kHz were downsampled to 100 Hz using an 8-pole bessel filter. Estimation of current jumps were carried out using a custom script with appropriate thresholds. Current jumps larger than 2 times the smallest observed jump were discarded for single channel histogram calculations for each design.

The current vs voltage readings for the three designs shown in Figure S13 were obtained using a step voltage ramping method. The applied voltage was toggled between positive and negative values starting from 25 mV while ramping the intensity by 25 mV during each positive cycle. The currents were recorded for 5 seconds under each voltage change and their means were used for the final plot. Measurements from three independent voltage ramping experiments were combined to get the final data points for fitting and conductance estimation. For TMB10_165, the voltages were only varied within the values of 150 and 75 for both positive and negative cycles because the currents at lower voltages were too close to the baseline and difficult to distinguish from baseline noise in this experimental setup.

#### Crystallography and structure determination of TMB10_163

SEC purified sample was used for crystallization. The crystallization screening was performed using a Mosquito LCP by STP Labtech. Crystals grew successfully in 3.25 M 1,6 hexanediol and 0.01 M HEPES pH 7.5. Crystals were harvested directly from a screening tray, and flash frozen in liquid nitrogen. X-ray diffraction was performed at ALS beamline 8.2.1, data were processed with XDS (*1*), and merged/scaled using Pointless/Aimless in the CCP4 program suite (*2*). The structure was phased by molecular replacement using the designed structure as the search model by Phaser (*3*) and refined with Phenix (*4*). Following molecular replacement, the models were improved, and efforts were made to reduce model bias. Structures were refined in Phenix. Model building was performed using COOT (*5*). The final model was evaluated using MolProbity (*6*). Data collection and refinement statistics are recorded in Table S1. Data deposition, atomic coordinates, and structure factors reported in this paper have been deposited in the Protein Data Bank (PDB), http://www.rcsb.org/ with accession code 8UZL.

##### TMB12_3 Expression for NMR

BL21(DE3) Lemo cells were transformed with a pET29b-derived expression plasmid for TMB12_3. Cells were grown in M9 minimal medium and expression was induced with 0.5 mM IPTG for 22h at 24°C. For expression of [*U*-99% ^2^H, ^15^N, ^13^C]-labeled samples, M9 was prepared with D_2_O, ^15^NH_4_Cl and deuterated-^13^C-glucose. For the expression of the [*U*-99%-^2^H, ^15^N] labeled sample, M9 was prepared with D_2_O and ^15^NH_4_Cl. For the selectively labeled [^15^N-Lys] and [^15^N-Phe] samples, the desired ^15^N labeled amino acid was added to the culture 45 min before induction.

##### Purification and Refolding of TMB12_3

Cells were lysed using a M110L from Microfluidics. Inclusion bodies were isolated and dissolved in denaturing buffer (20 mM Tris/HCl pH 8, 150 mM NaCl, 6 M GuHCl), then dialyzed against H_2_O in a 10,000 MWCO dialysis membrane for 2 h, followed by centrifugation at 30,000 g to precipitate the protein. Precipitated TMB12_3 was dissolved in 10 mM Tris/HCl pH 8, 7 M urea. Refolding was performed at 4°C by dropwise rapid dilution into a stirred refolding buffer (20 mM Tris/HCl, 2 mM EDTA, 0.6 M L-Arg, 15 mM LDAO, pH 10). The dilution ratio was set to 1:20 and after overnight stirring, the refolded protein was dialyzed against 20 mM sodium phosphate buffer pH 6.8, 1 mM EDTA for 2h. The refolded protein was concentrated with MWCO 10,000 and the sample was loaded on an S200 size exclusion column, pre-equilibrated with 20 mM sodium phosphate buffer, 1 mM EDTA, 15 mM LDAO. The fractions containing protein were pooled and concentrated using MWCO 10,000.

##### Isotope labeling samples

The following samples were made: [U-99%-^15^N]-TMB12_3 in LDAO, [U-99%-^2^H, ^15^N, ^13^C]-TMB12_3 in LDAO, [U-99%-^2^H,^15^N; 99%-^1^H^β^,^13^C^β^-A; 99%-^1^H^ε^,^13^C^ε^-M; 99%-^1^H^δ1^,^13^C^δ1^-L; 99%-^1^H^ɣ1^,^13^C^ɣ1^-V]-TMB12_3 in [U-99%-^2^H]-LDAO, [^15^N-Lys]-TMB12_3 in LDAO, [^15^N-Phe]-TMB12_3 in LDAO. The final sample conditions were 20 mM sodium phosphate buffer, 1 mM EDTA, pH 6.8, 300–500 mM LDAO, 0.2–1 mM TMB12_3.

##### NMR experiments

All experiments were carried out at 25°C on Bruker spectrometers operating at field strengths of 700, 800 and 900 MHz. All spectrometers were equipped with a cryogenic triple-resonance probe. The following experiments were recorded: 2D [^15^N, ^1^H]-BEST-TROSY (*7*), 3D BEST-TROSY-HNCACB with ^2^H decoupling (*7*), 3D [^1^H,^1^H]-NOESY-^15^N-TROSY (*8*), 2D ^13^C-Methyl-SOFAST (*7*), 3D [^1^H,^1^H]-NOESY-^13^C-HMQC (*9*).

##### Structure calculation

Structure calculation was performed with CYANA 3.98.15 (*10*). All spectra were processed and analyzed with NMRPipe (*11*) and ccpNMR version 3 (*12*). 129 dihedral constraints were derived by TALOS-N (*13*) from the experimentally determined C_ɑ_, C_β_, N and HN chemical shifts. Only TALOS-N predictions classified as “strong” were used. Tolerances were set to one standard deviation, capped at a maximum of 20°. A total of 213 experimental NOEs were obtained from 3D [^1^H,^1^H]-NOESY-^15^N-TROSY and 3D [^1^H,^1^H]-NOESY-^13^C-HMQC spectra. 48 Hydrogen bond constraints were inferred from the measured NOE data with upper limits of 2.0 / 3.0 Å and lower limits of 1.8 / 2.7 Å for the HN…O and N…O, respectively. In regions with sparse assignment, these constraints were inferred indirectly from the experimentally established β-strand topology (dashed black lines in Figure 3F). A total of 400 structures were calculated and the ensemble of 20 lowest energy structures was selected. Ramachandran statistics of this ensemble showed 76.6% of residues in most favored regions, 20.0% in allowed regions, 2.6% in generously allowed regions and 0.8% in disallowed regions.

##### Calculating the theoretical diameters of nanopores

The diameters of the nanopores were inferred from the observed single-channel conductance (*G* (nS)) using the access resistance model (**eq. 1**), where R is the resistance, S the conductivity of the solution (S/m), d is the diameter of the pore (nm) and L is the pore length (nm). The purely geometric model approximates the properties of a cylindrical nanopore and assumes homogeneous solution, pore and membrane neutrality and constant potential at the pore mouth.

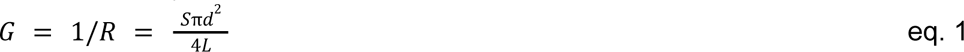

A pore length of 3.5 nm was used for all the calculations based on the total transmembrane span of the TMB designs. The conductivity of a solution of 0.5 M NaCl was estimated to be 40.5 mS/cm based on the previously reported relationship between NaCl molarity and conductivity of the solution.

##### Folding kinetics measured by tryptophan fluorescence

Protein samples were buffer exchanged into unfolding buffer (50 mM glycine-NaOH pH 9.5, 8 M/10 M urea) using a 0.5 mL ZebaSpin 7K MWCO desalting column. The concentration was determined by nanodrop. For kinetic experiments, 15 µL of unfolded protein was added to 485 µL of pre-warmed (25 °C) pre-fold buffer of LUVs in 50 mM glycine-NaOH, pH 9.5, in a QS quartz cuvette to give final concentrations of 0.4 µM OMP, 600-3200 LPR (*mol:mol*), 0.24-4 M urea, 50 mM glycine-NaOH pH 9.5. Immediately after mixing, a time based fluorescence scan was carried out on a PTI QuantaMaster™ spectrofluorometer (Photon Technology International), controlled by FelixGX v4.3 software, with excitation at 280 nm, and emission measured at 335 nm. The slit settings were 0.5 nm for excitation, and 5 nm for emission, to minimize photobleaching. Integration was set at 1 s between time points, and the temperature was maintained at 25 °C throughout. Fluorescence emission spectra were measured by exciting tryptophan at 280 nm, and measuring fluorescence emission between 300-400 nm, using the same slit-width settings as above, with samples in urea concentrations between 0.24-9.9 M urea.

##### Circular dichroism in liposomes

Protein samples were prepared in a similar manner as for the tryptophan fluorescence samples. Samples were made to a 600:1 (*mol:mol*) LPR, with final concentrations of 4 µM TMB, 1.2 mM lipid-LUV, 0.24-8 M urea, 50 mM glycine-NaOH pH 9.5 in a final reaction volume of 300 µL. The reaction was allowed to proceed overnight at 25 °C to maximize the fraction of protein folded into the lipid-LUV bilayer. Controls were made where the volume of substrate was replaced with 50 mM glycine-NaOH pH 9.5 and an appropriate volume of urea to match the protein samples. These were used to normalize the data by subtracting their CD signals from the CD signal from the protein containing samples. Measurements were taken using 300 µL of sample in a 1 mm QS quartz cuvette, using a Chirascan plus CD Spectrometer (Applied Photophysics). The bandwidth was set at 2.5 nm, and used adaptive sampling to adjust the integration time for the optimal signal:noise. Four scans were averaged between 260 nm to the lowest useable wavelength for each respective sample, which was the point where the voltage reached its upper limit of 1000 V, after which the data became unusable. During temperature ramp experiments, only single scans were taken as the temperature ranged between 25 °C and 87 °C.

##### Equilibrium denaturation analysis

To determine the urea dependence of TMB folding, urea denatured TMBs in 50 mM glycine-NaOH pH 9.5, 10 M urea were diluted into DUPC LUVs at an Lipid-to-Protein ratio (LPR) of 600:1 (mol/mol) to give a final concentration of 0.4 µM TMB in 50 mM glycine-NaOH pH 9.5 containing 2-9.9 M urea, and folding was allowed to proceed overnight at 25°C. For urea dependence of unfolding, TMBs were folded in DUPC LUVs (LPR 600:1 (mol/mol)) in 50 mM glycine-NaOH pH 9.5, 2 M urea overnight at 25°C. Pre-folded TMBs were then unfolded by dilution into 50 mM glycine-NaOH pH 9.5 containing 2-9 M urea to a final TMB concentration of 0.4 µM and incubated overnight at 25 °C. Tryptophan fluorescence emission spectra were obtained using a PTI QuantaMaster spectrofluorometer (Photon Technology International) in QS quartz cuvettes with excitation slits set to 1 nm and emission slits set to 5 nm. Fluorescence was excited at 280 nm and emission spectra were acquired between 300-400 nm using a step size of 1 nm and an integration time 0.5 seconds. The average wavelength between 325-375 nm was calculated using equation 2, where <λ> is the average wavelength, I_λ_ is the fluorescence intensity at a given wavelength, λ is the wavelength, and ∑I is the sum of the intensity of the entire emission spectra.

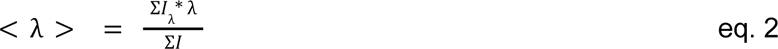

The experimental data were fitted to a 2-state transition model(*14*) to extract ΔG_0_ (the Gibb’s free energy for unfolding in the absence of denaturant), the m-value (m_UF_), the global dependence of ΔG_0_ on the concentration of denaturant and C_m_ (the transition midpoint) based on Obs_F_ and Obs_U_ (the observed <λ> for the folded and unfolded states in the absence of denaturant ([D]=0)), m_f_ and m_u_ (the linear dependence of Obs_F_ and Obs_U_ to [D]). The observed <λ> was corrected to account for the difference in quantum yield between the folded and unfolded states based on the Q-factor (QF), which is calculated by taking the ratio of the summed fluorescence intensities at the folded and unfolded states. R is the universal gas constant and T is the absolute temperature.

##### *De novo* backbone assembly

The Rosetta blueprint representations of the beta-barrel backbones were generated based on user input using a custom python script available on GitHub https://github.com/vorobieva/demo_TMB_design/tree/master/generate_blueprint. The script requires the SciPy and BioPython modules and generates a Rosetta blueprint (describing local secondary structure and torsion angle bins per residue) and a Rosetta constraints file (describing backbone-backbone hydrogen bonds in the barrel). Examples of blueprint and constraints files used in this study are available on GitHub https://github.com/vorobieva/demo_TMB_design/tree/master/12_strands_square/assemble_backbones. The backbones were assembled based on such a blueprint and constraints file using the Rosetta BluePrintBDR application (*15*), alternating between sampling of backbone fragments and minimization with hydrogen bonds constraints. The highest-quality protein backbones (250-500 backbones, based on Rosetta `vdw`, `omega` and `rama_prepro` scores) were selected as template for combinatorial sequence design.

##### Combinatorial TMB sequence design

*De novo* β-barrel backbones assembled using the Rosetta coarse-grained centroid model were subject to one round of fast atomistic refinement of the backbones with the Rosetta full-atom model (`ref2015` energy function with limited sampling depth `-nstruct 1` and limited sequence space). The Tyr-Gly-Asp/Glu TMB folding motifs were then designed into the structures using the HBNet (*16*) Rosetta application, which finds all possible positions of the hydrogen bond acceptor residue in the motif (Asp or Glu) based on defined tyrosine positions. The refined TMB backbones were used as templates for several rounds of combinatorial sequence design, alternating between the water-accessible pore (two rounds of design) and the lipid-exposed surface (three rounds of designs) using re-fitted energy functions (see below). Rosetta resfiles were used to define the set of amino acids sampled at each position in the protein. The surface-exposed residues were constrained to mostly hydrophobic amino acids while all amino acids but PRO and CYS were allowed at pore-lining positions. The β-turn residues were designed using previously identified canonical TMB β-turn sequences (Figure S2). At each iteration of sequence optimization, the whole population of designs was analyzed, population-wide selection metrics were computed (see below) and around 10 % of the designs were selected to be used as input for the next iteration. After an iteration of pore-residue design, the outputs were selected based on backbone quality metrics (Rosetta `omega`, `rama_prepro`, `hbond_lr_bb` scores) and on the computed `total` and `hbond_sc` energies of the hydrogen bond acceptor residues in the Tyr-Gly-Asp/Glu folding motifs. After an iteration of surface-exposed residue design, the outputs were selected based on Rosetta `total_energy` and on the retention of the Tyr-Asp/Glu interactions in the designed folding motifs (which are repacked while the surface residues are designed). The resfiles used in this study to generate the different TMB architectures are available on GitHub (https://github.com/vorobieva/demo_TMB_design/tree/master). A complete description of the design pipeline, analysis scripts and example inputs are available on GitHub (https://github.com/vorobieva/demo_TMB_design/tree/master).

##### Re-fitted TMB-specific energy functions

To fine-tune the amino acid propensities to TMB-specific statistics, the Rosetta reference energy function (`ref2015`) was modified by testing several variations of weights on one representative TMB backbone. To generate an energy function to design the water-accessible pore-lining residues, the weight of the Rosetta full-atom solvation energy (`fasol`), electrostatic energy (`faelec`) and of the reference energies of small disorder-promoting amino acids (ALA and SER) were systematically varied. The scoring function used for subsequent sequence design was selected based on the closest match between the resulting designed sequences and naturally-occurring TMB sequences at the level of the overall hydropathy of the pore and the frequency of ALA and SER amino acids. To generate an energy function to design the lipid-exposed surface residues, the weight of the Rosetta full-atom solvation energy and of the reference energies of large hydrophobic (PHE) and of small disorder-promoting amino acids (GLY and ALA) were systematically varied. A scoring function was selected based on the closest match between the resulting designed sequences and naturally-occurring TMB sequences at the level of the overall hydropathy of the surface and the frequency of ALA and GLY amino acids.

Comparison of the scoring weights optimized to design the pore residues of TMB12s designs.

|  | TMB12_square | TMB12_oval | TMB12_rectangle | ref2015 |
| --- | --- | --- | --- | --- |
| `fasol` | 0.9 | 0.9 | 0.9 | 1.0 |
| `faelec` | 1.4 | 1.45 | 1.45 | 1.0 |
| ALA `refE` | 0.4 | -0.6 | -0.6 | 1.32468 |
| SER `refE` | -1.2 | -1.8 | -1.8 | -0.28969 |

##### Designs validation and selection

The final designs were filtered based on the desired balance between local secondary structure frustration (β-sheet propensity of the sequence between 30 % and 50 % (predicted with RaptorX (*17*)) and aggregation propensity score predicted with Tango (*18*) smaller than 1500) and sequence-encoded tertiary structure. The sequence/structure compatibility was assessed based on the capacity of AlphaFold2 (*19*) to fold the sequence into the designed TMB structure in single sequence mode (no multiple sequence align input) and using 48 recycles through the network. Selected predictions had AlphaFold2 plDDT scores higher than 0.8 and showed high structure similar to the expected design model when superimposed with TMAlign (*20*), with calculated Root Mean Square Deviations (RMSD) of less than 2.1 Å.

### Supplementary Table

**Table S1.** X-ray diffraction data collection and refinement statistics.

|  | TMB10_163 (8UZL) |
| --- | --- |
| Resolution range | 49.92 - 2.5 (2.58 - 2.5) |
| Space group | $P\ 2_1$ |
| Unit cell | 55.62, 52.55, 100.05, 90, 93.67, 90 |
| Unique reflections | 20045 (1961) |
| Multiplicity | 4.3 (4.5) |
| Completeness (%) | 98.87 (98.54) |
| Mean I/sigma (I) | 4.26 (0.36) |
| Wilson B-factor | 43.07 |
| R-merge | 0.1255 (0.9234) |
| R-pim | 0.06884 (0.4931) |
| CC<sub>1/2</sub> | 0.99 (0.832) |
| Reflections used in refinement | 20021 (1960) |
| R-work | 0.2272 (0.3356) |
| R-free | 0.2624 (0.3828) |
| Number of non-hydrogen atoms | 4535 |
| macromolecules | 4420 |
| ligands | 16 |
| solvent | 99 |
| Protein residues | 568 |
| RMS (bonds) | 0.002 |
| RMS (angles) | 0.45 |
| Ramachandran favored (%) | 96.96 |
| Ramachandran allowed (%) | 3.04 |
| Ramachandran outliers (%) | 0.00 |
| Average B-factor | 52.26 |
| macromolecules | 52.31 |
| ligands | 52.13 |
| solvent | 50.48 |

Statistics for the highest-resolution shell are shown in parentheses.

**Table S2.** NMR and refinement statistics for TMB12_3 in LDAO micelles.

|  | TMB12_3 |
| --- | --- |
| NMR distance and dihedral constraints | 442 |
| Distance constraints | 313 |
| Total NOE | 213 |
| Intra-residue | 22 |
| Inter-residue | 191 |
| Sequential ($\vert i - j\vert = 1$) | 80 |
| Medium-range ($\vert i - j\vert < 4$) | 24 |
| Long-range ($\vert i - j\vert > 5$) | 109 |
| Intermolecular | 0 |
| Hydrogen bonds | 100 |
| Total dihedral angle restraints | 129 |
| f | 63 |
| y | 66 |
| Structure statistics |  |
| Violations (mean and s.d.) |  |
| Distance constraints (Å) | 0.02 +/- 0.07 |
| Dihedral angle constraints (°) | 2.6 +/- 1.3 |
| Max. dihedral angle violation (°) | 19 +/- 12 |
| Max. distance constraint violation (Å) | 0.46 +/- 0.21 |
| Deviations from idealized geometry | 0 |
| Average pairwise r.m.s. deviation* (Å) |  |
| Heavy | 3.23 |
| Backbone | 2.54 |

\* Calculated among 20 refined structures

**Table S3.**
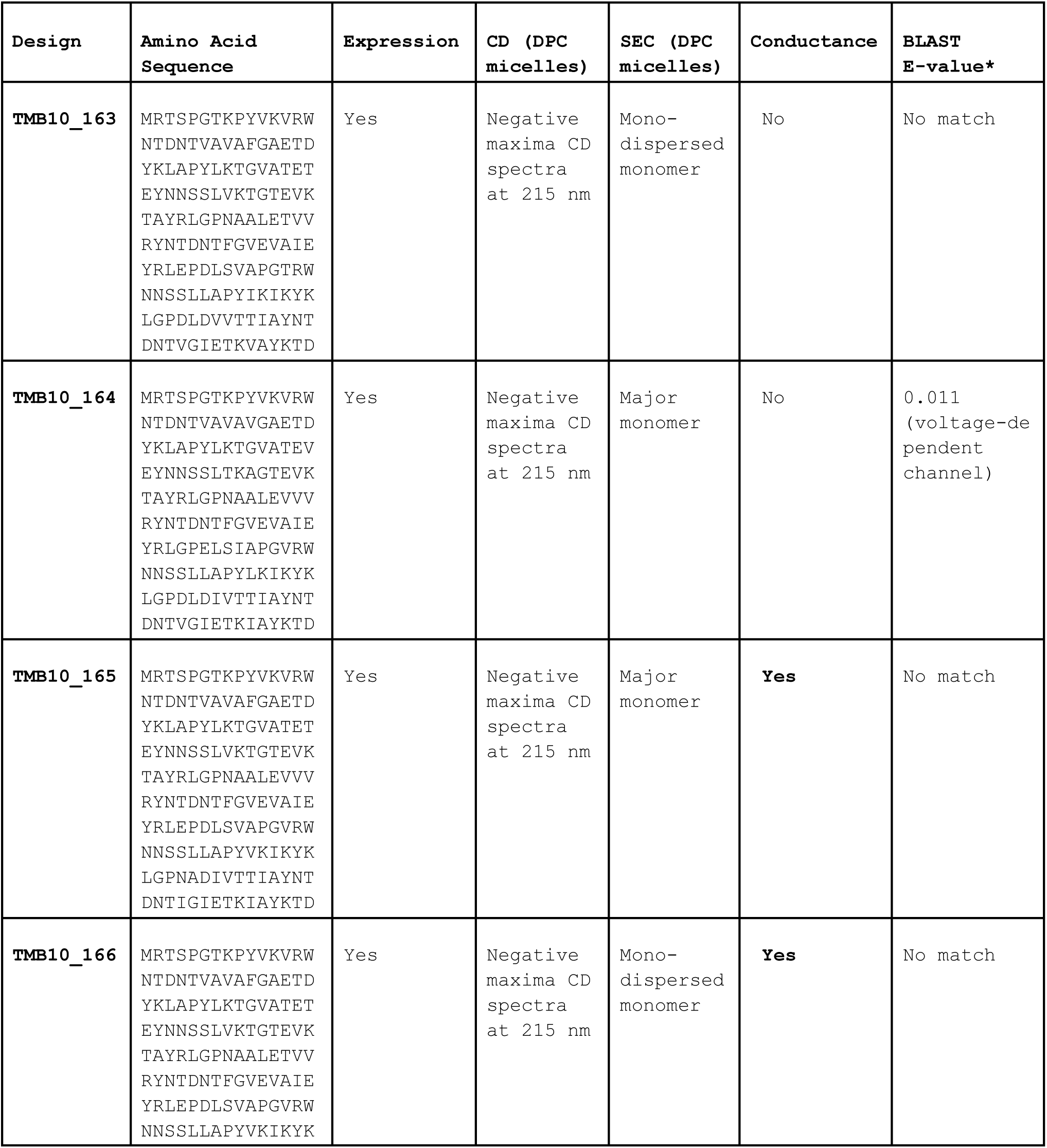
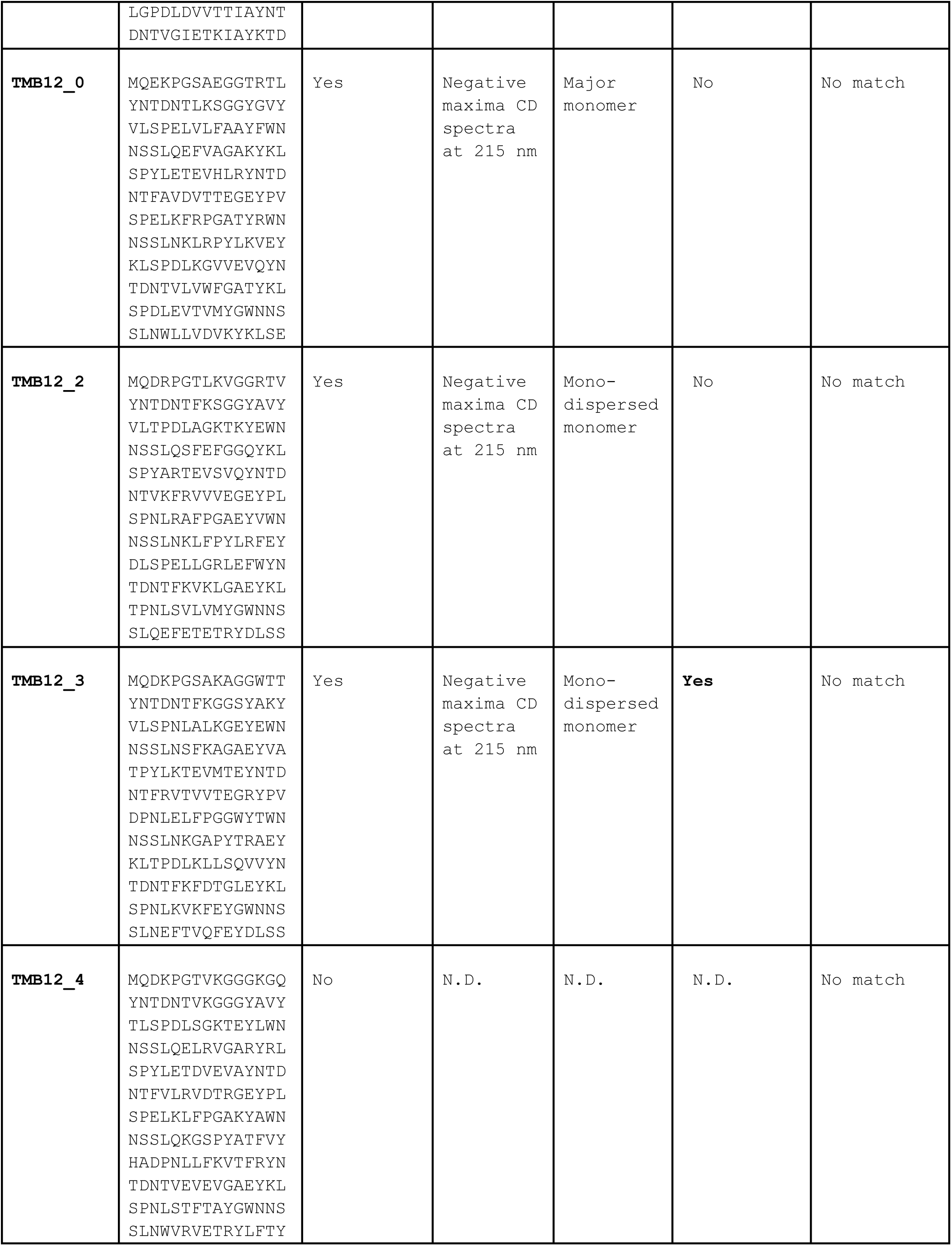
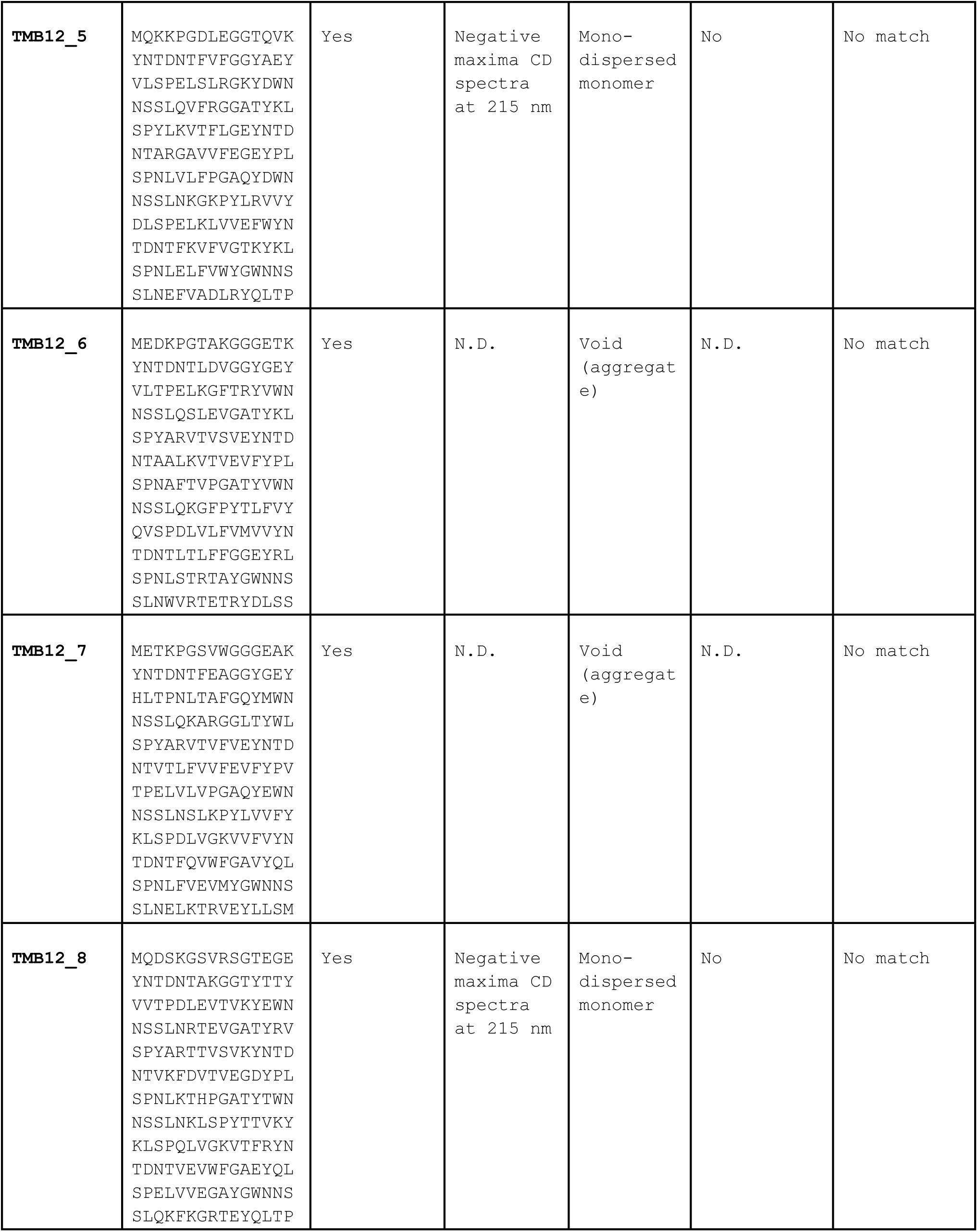
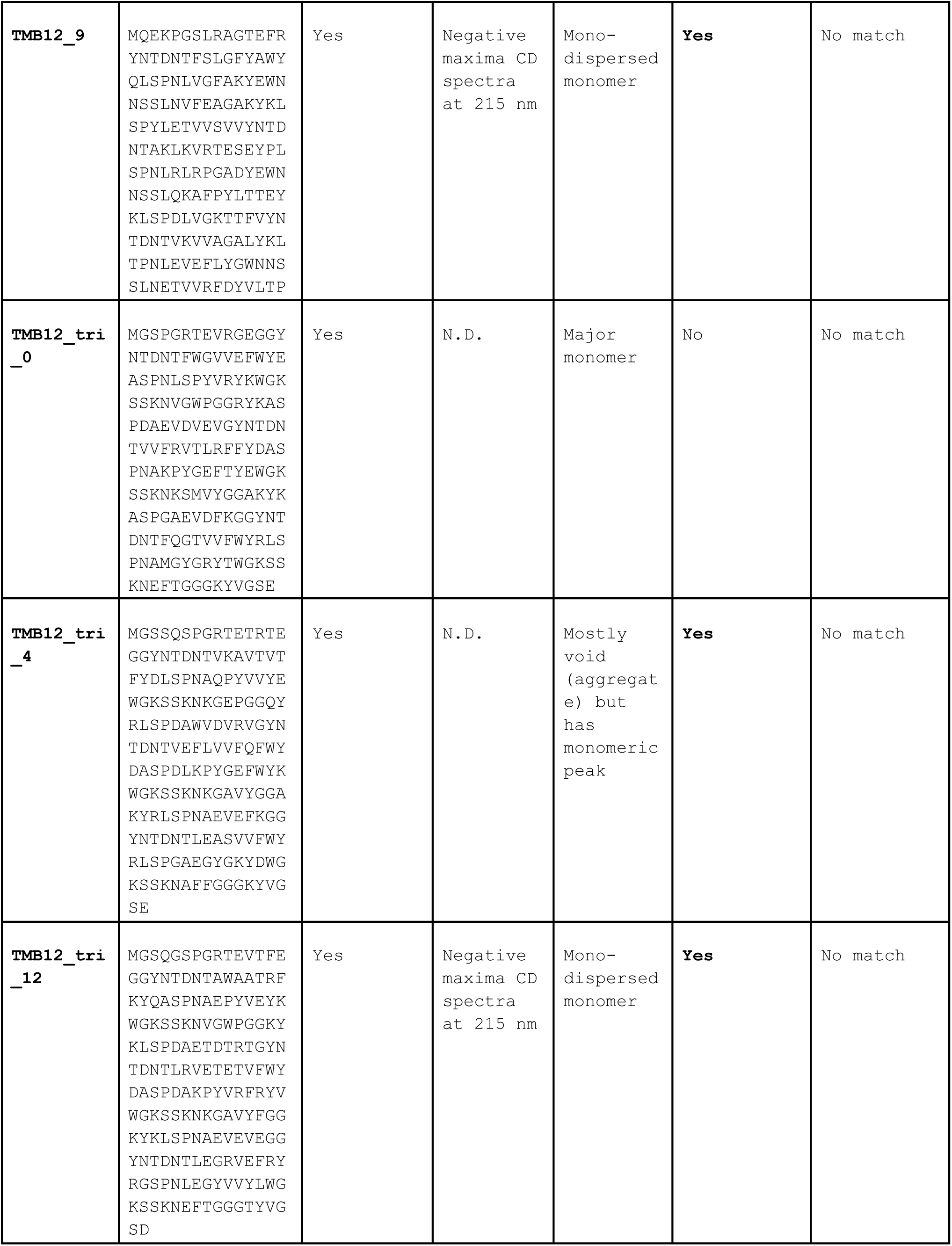
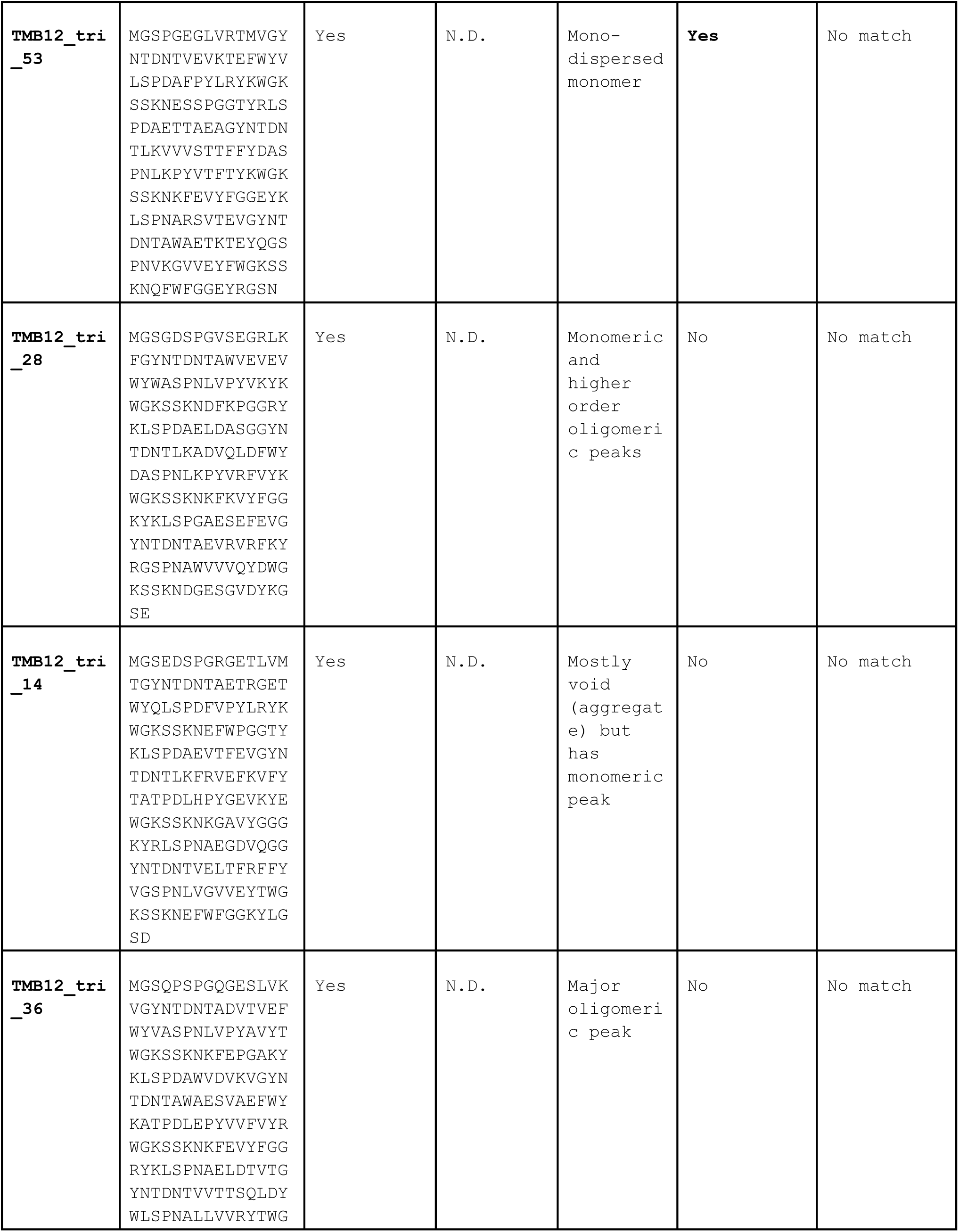
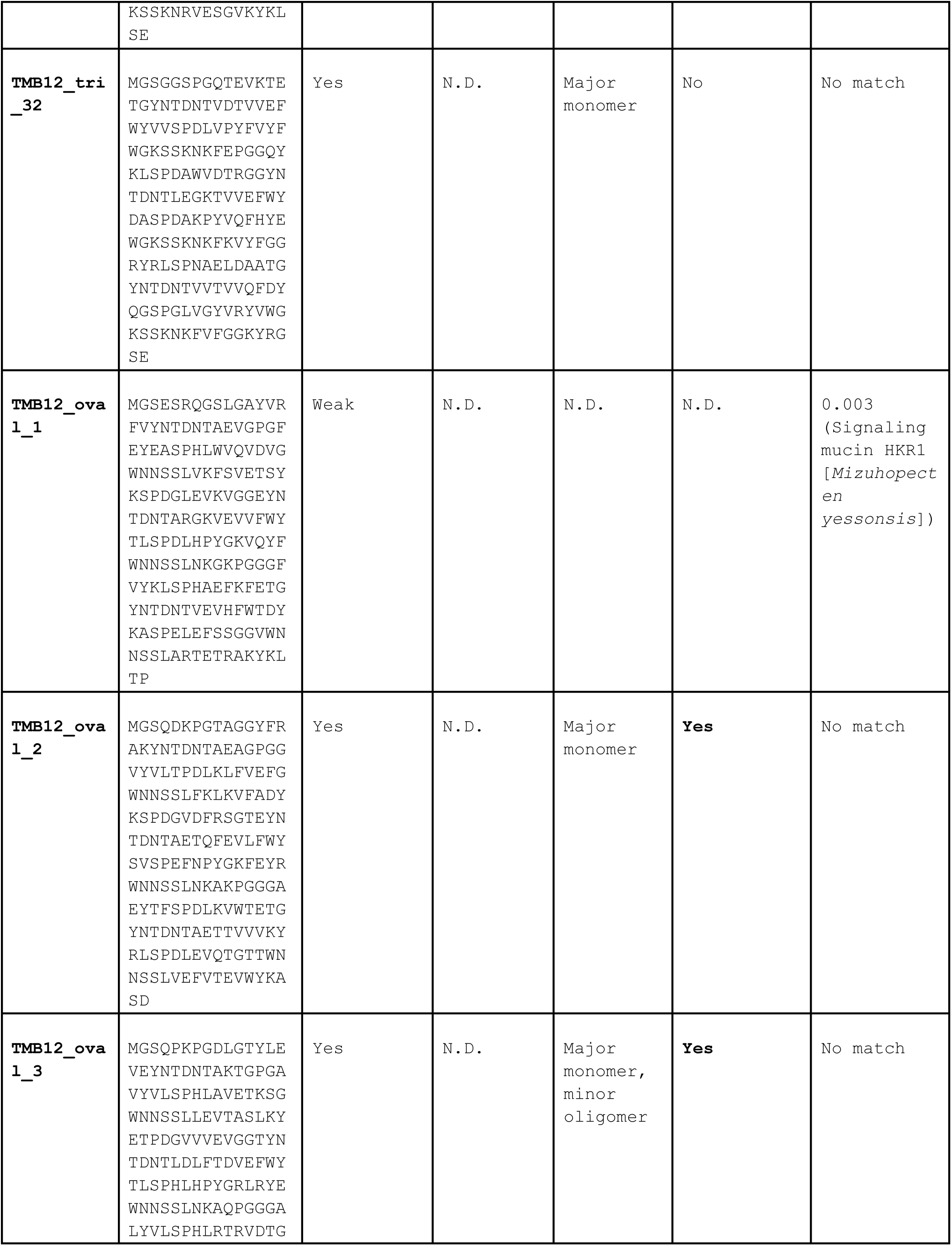
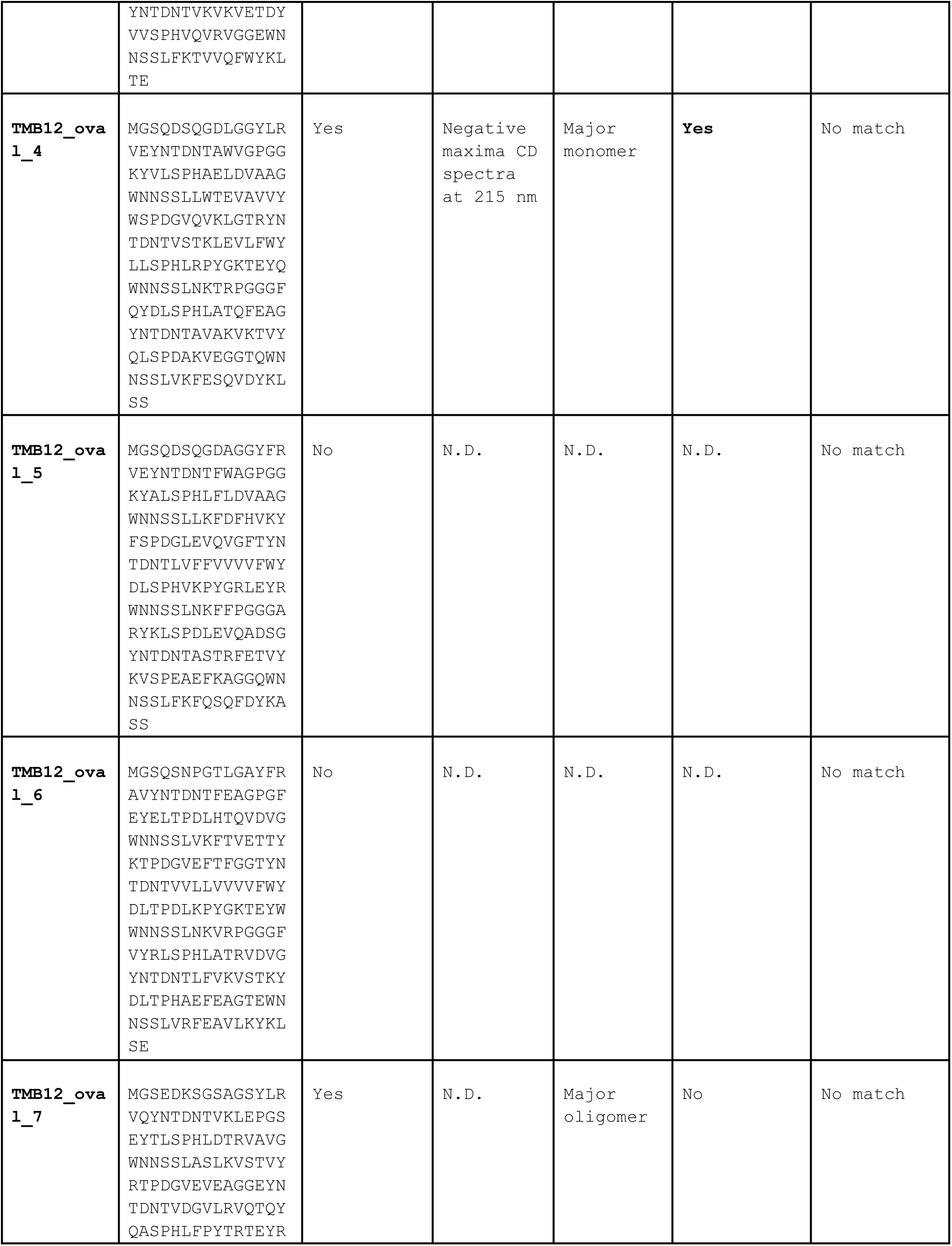
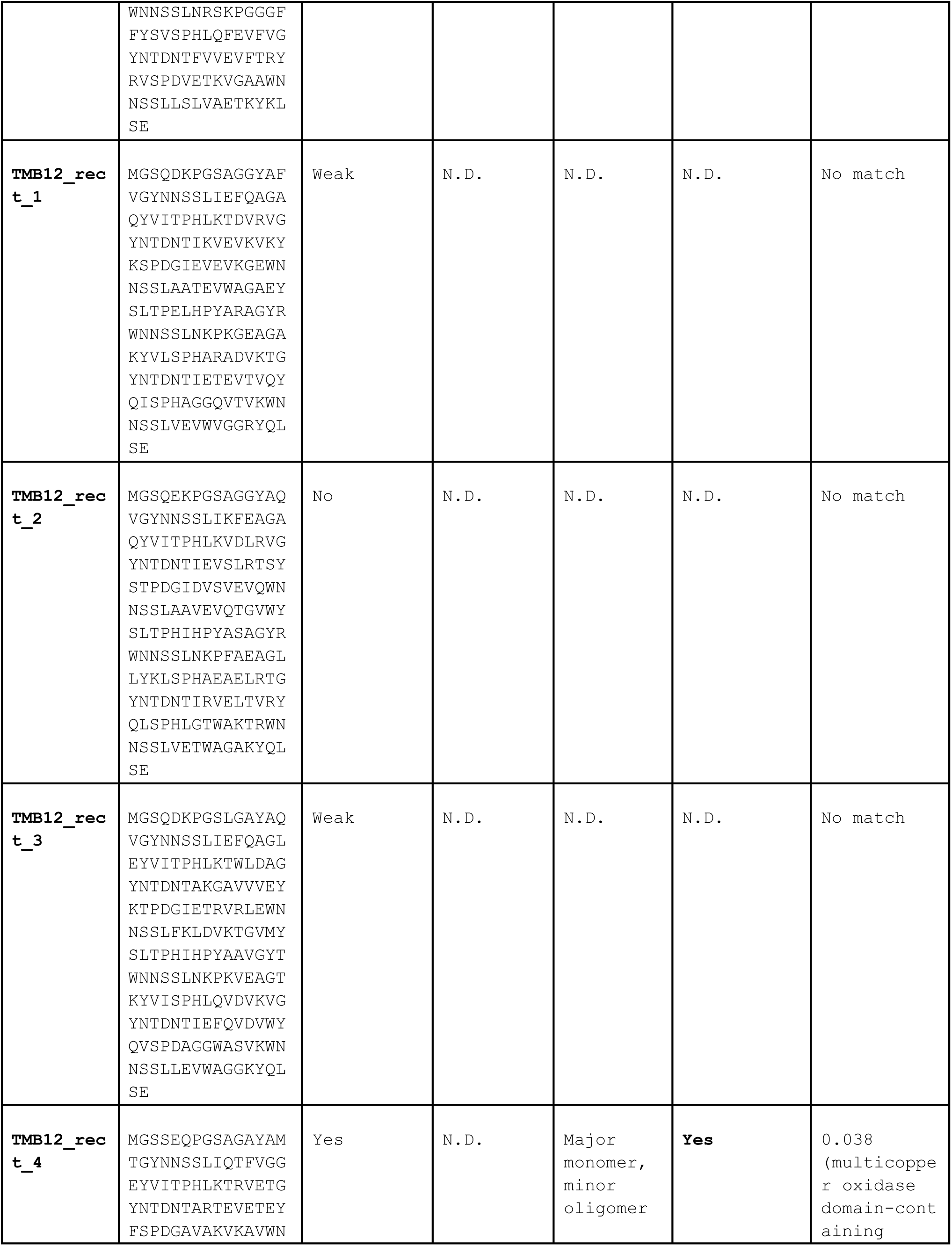
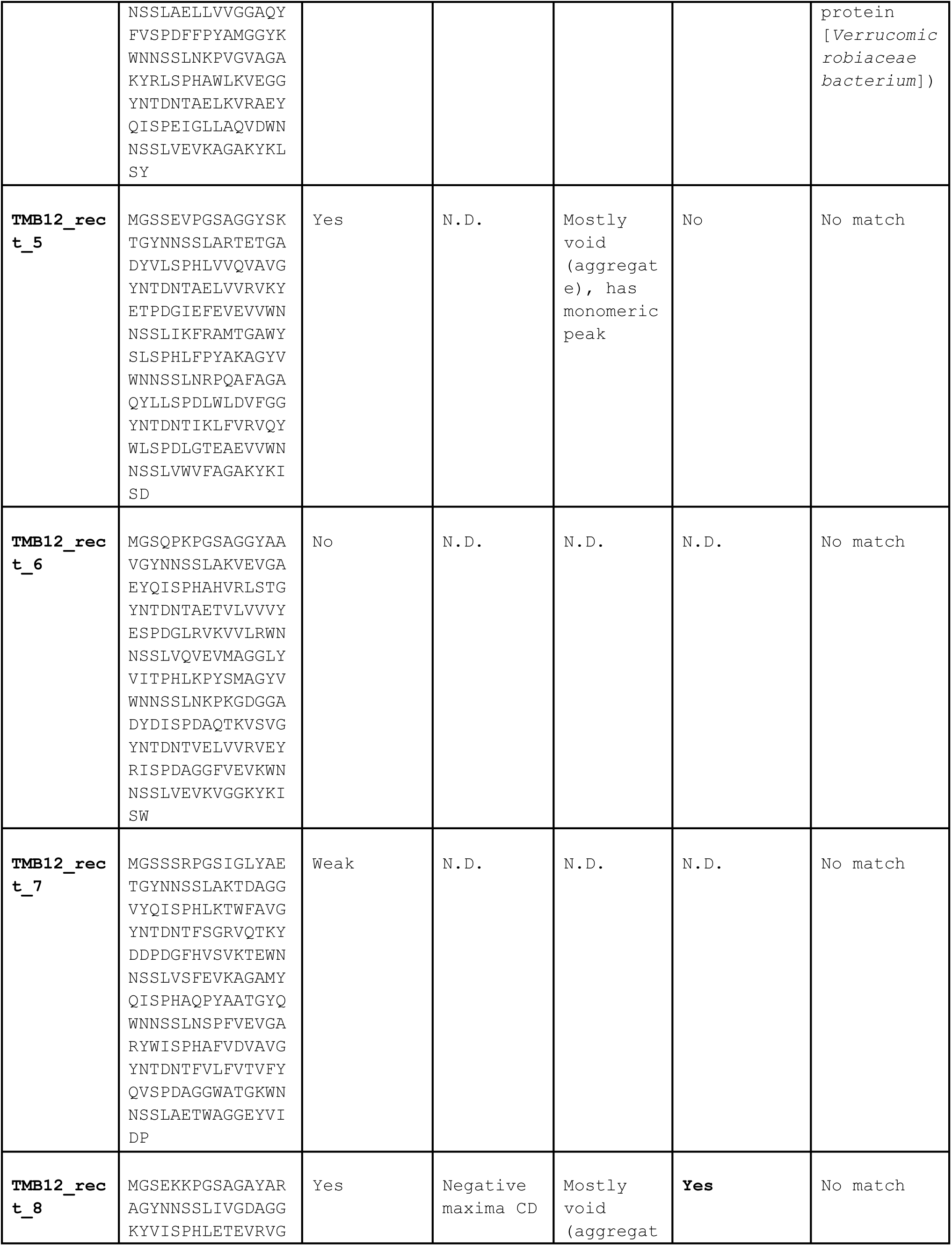
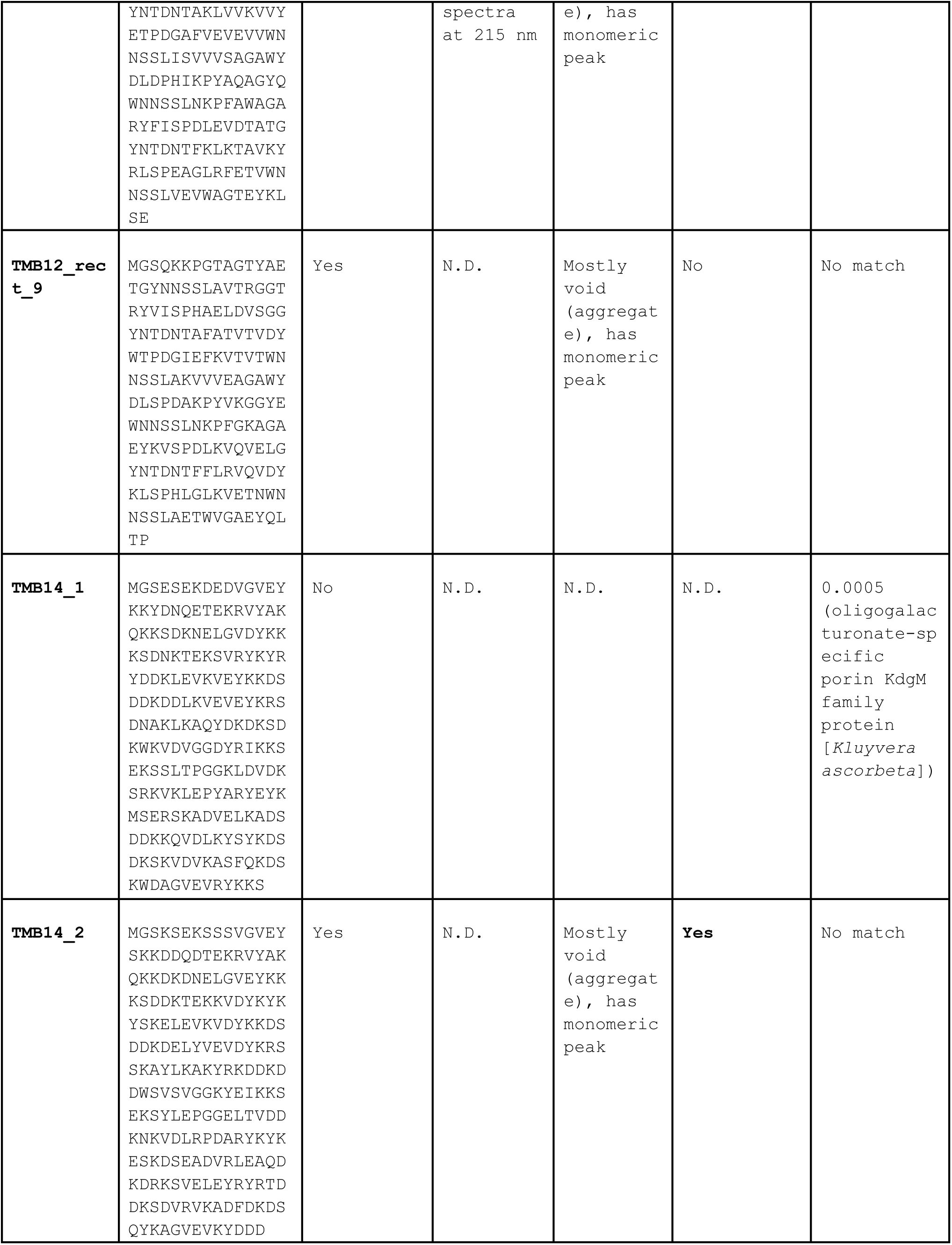
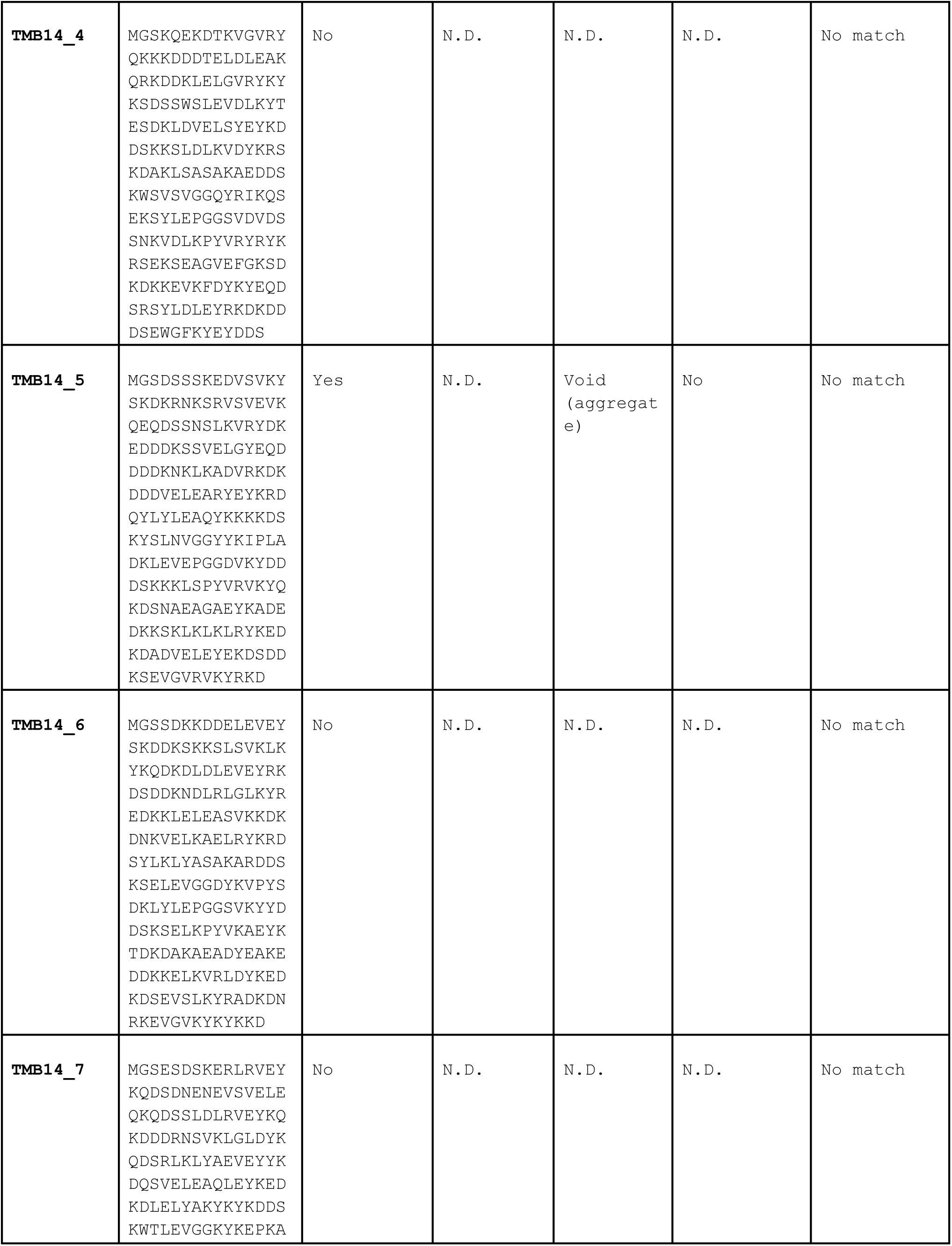
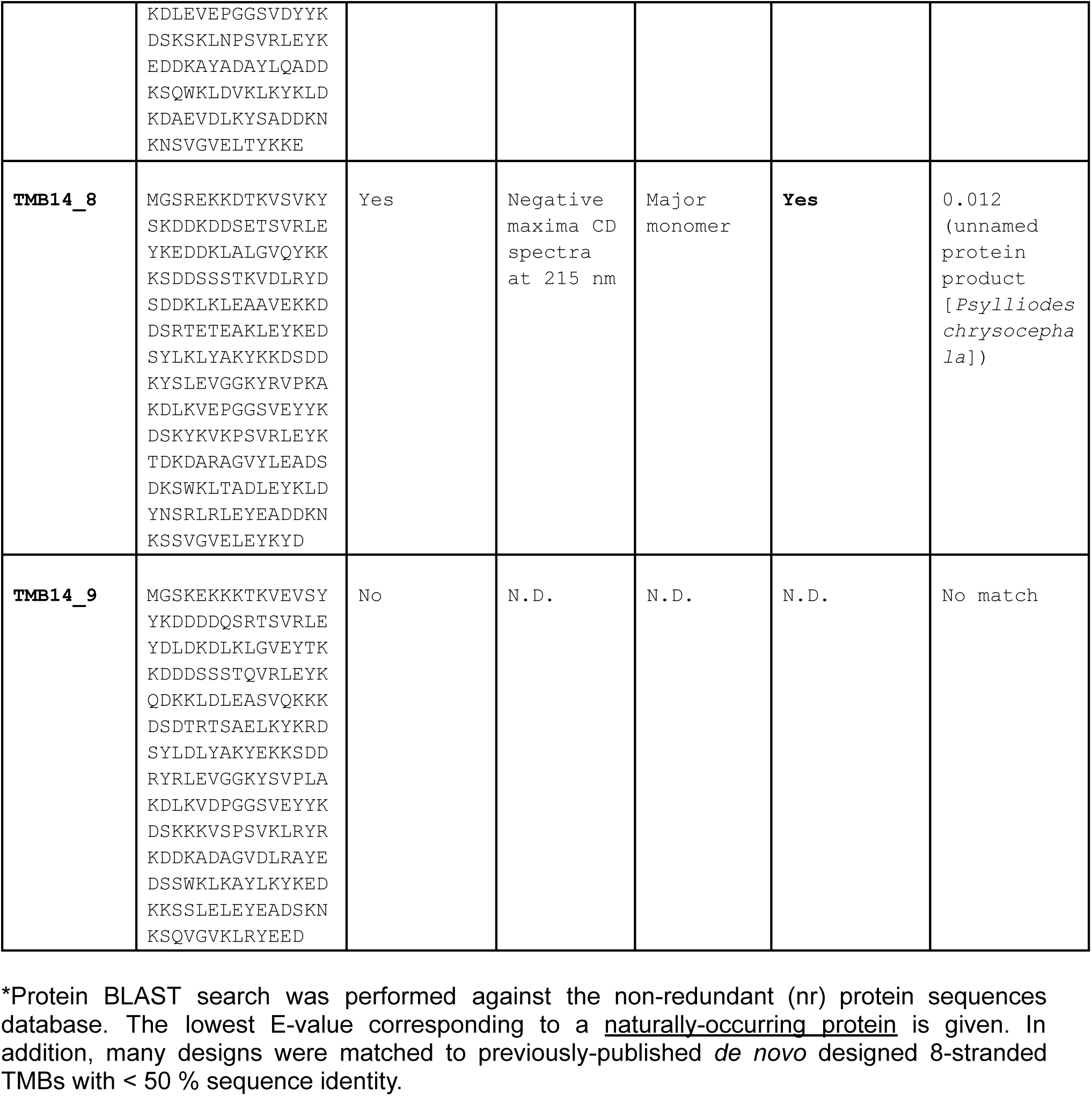
Experimentally tested designs.

### Supplementary Figures

**Figure S1:**
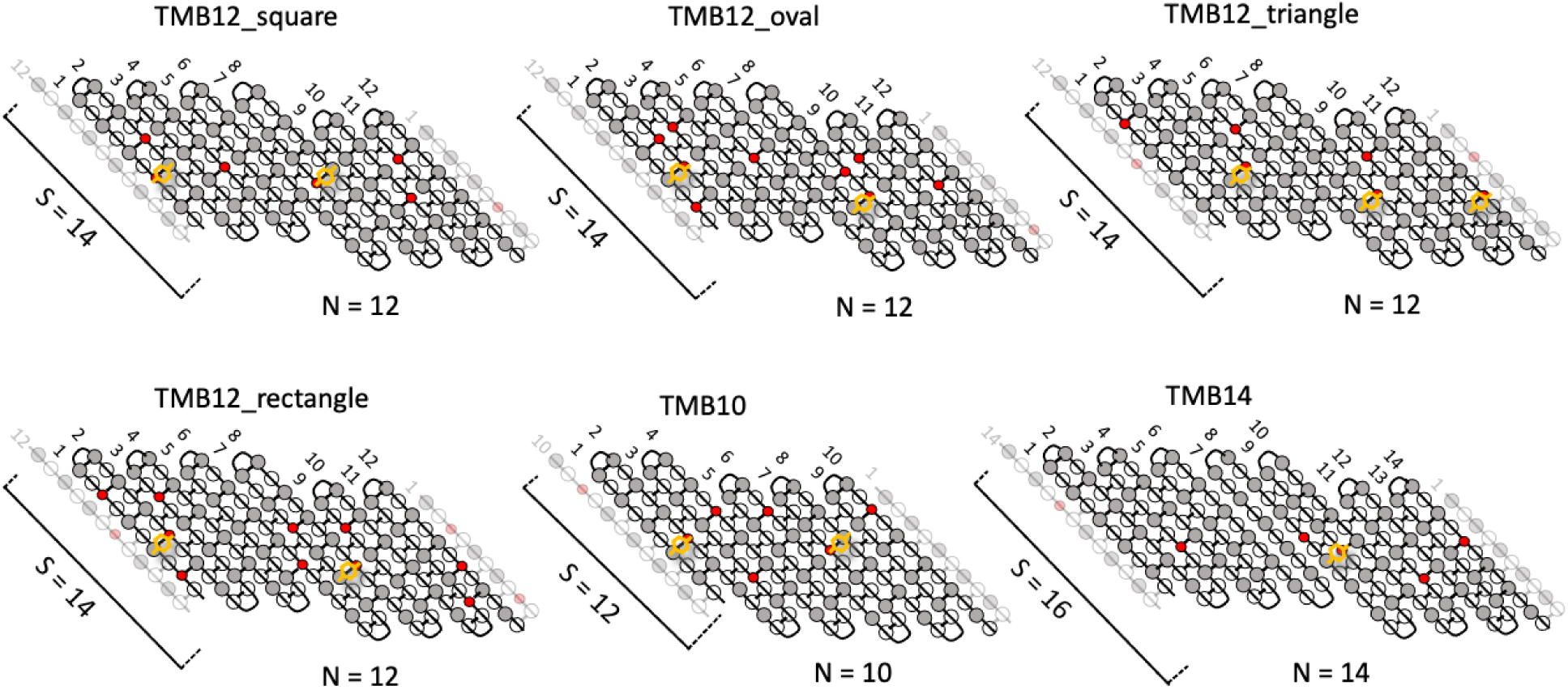
Six new β-barrel blueprints were generated here: one β-barrel of 10 strands (N=10) and a shear number of 12 (S=12), one β-barrel of 14 strands (N=14) and a shear number of 16 (S=16) and three β-barrels of 12 strands (N=12) and a shear number of 14 (S=14). The residues facing the β-barrel lumen and surface are shown as gray and white circles, respectively. Glycine kinks are shown as red circles and are facing the lumen. The tyrosine residues belonging to the Tyr-Gly-Asp/Glu folding motif are shown in orange.

**Figure S2:**
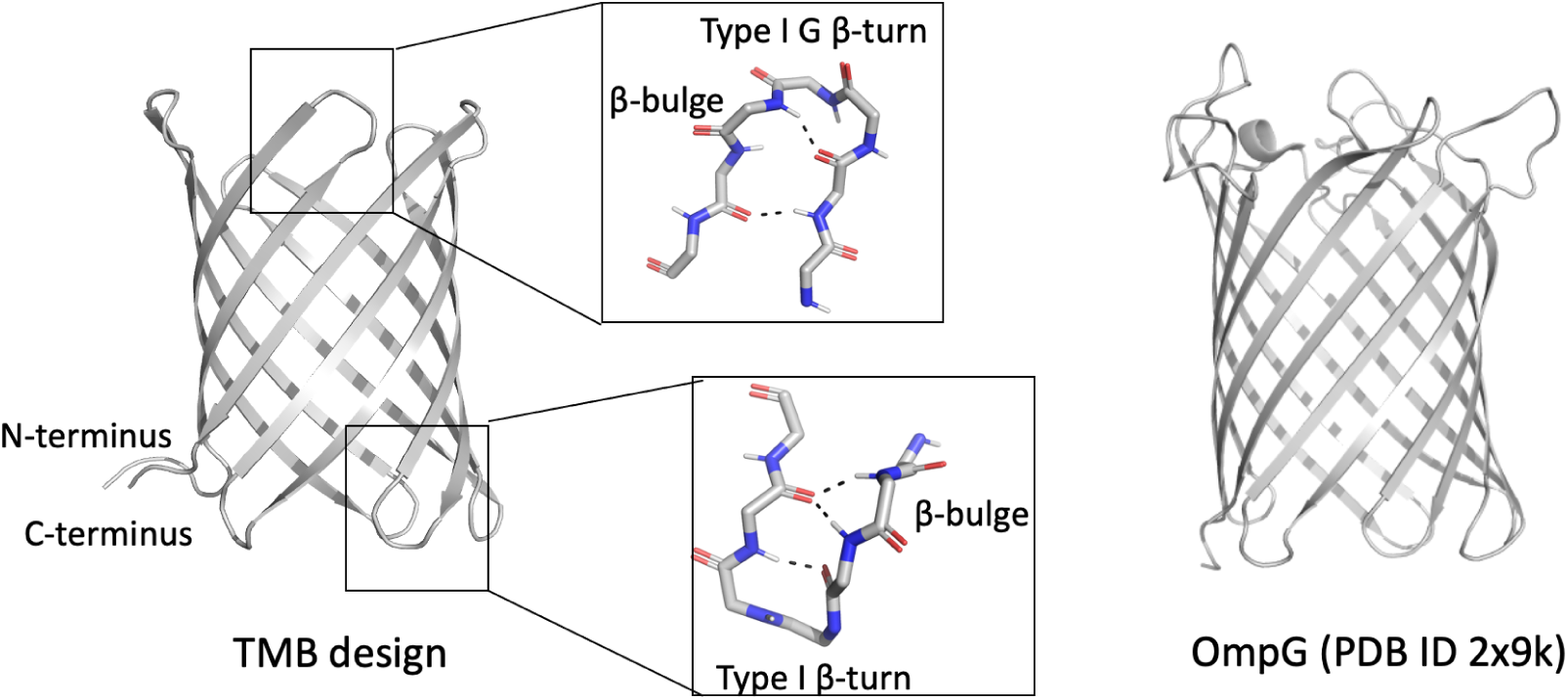
The β-strands of *de novo* designed TMBs are connected with short β-turns on both sides of the barrel: *cis*-hairpins (N- and C-termini side) are connected with canonical type I β-turns preceded by a β-bulge; *trans*-hairpins are connected with type I β-turns directly followed by a G-bulge. By comparison, naturally-occurring TMBs (exemplified here by OmpG, right) feature mostly long, disordered, loops on the *trans* side.

**Figure S3:**
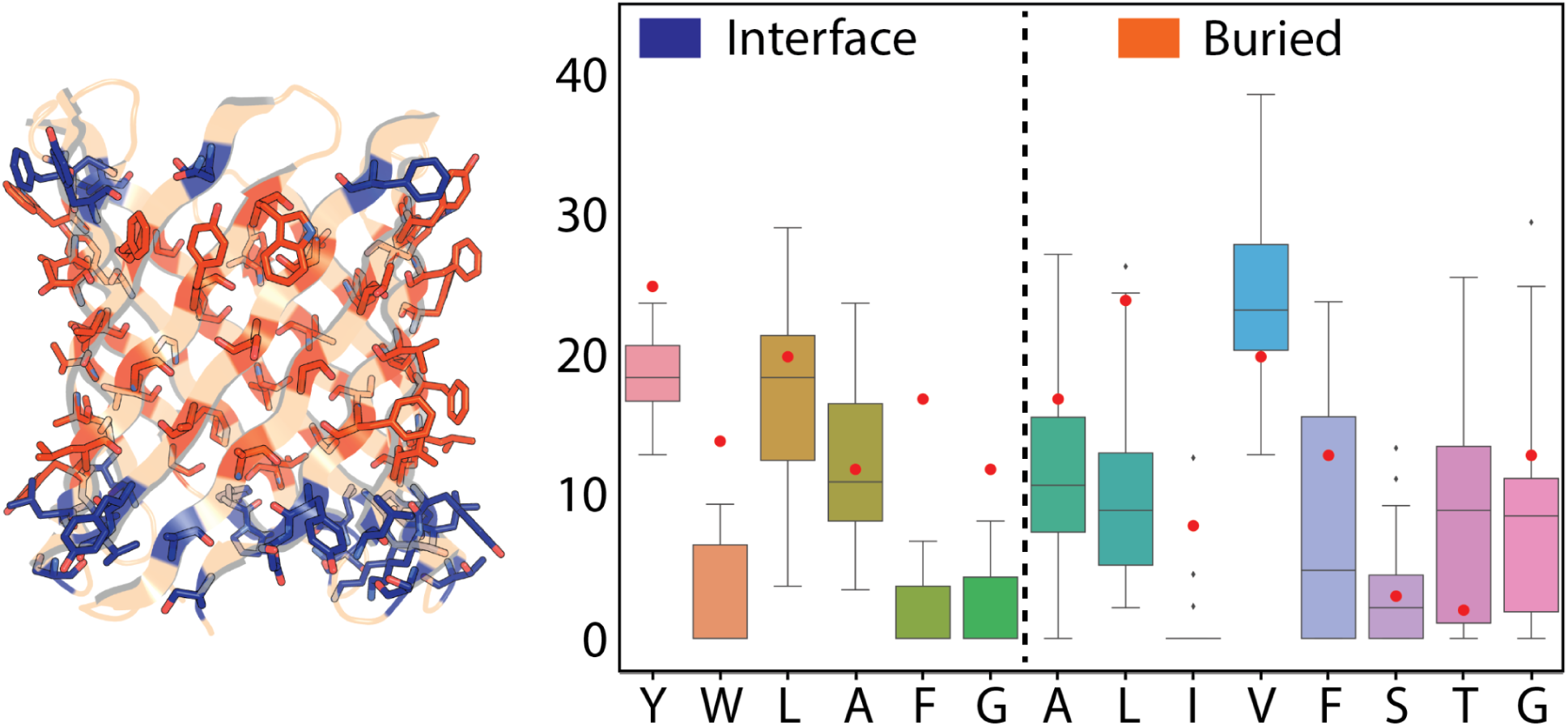
Amino acid composition of the membrane exposed surface of designed beta-barrel pores. Y-axis is calculated composition among all types of amino acids in the interface and buried region for each design respectively. The red dots are averaged amino acid compositions for the indicated amino acids in the respective regions over all transmembrane beta-barrel proteins in the OPM (Outer Membrane Protein) database.

**Figure S4:**
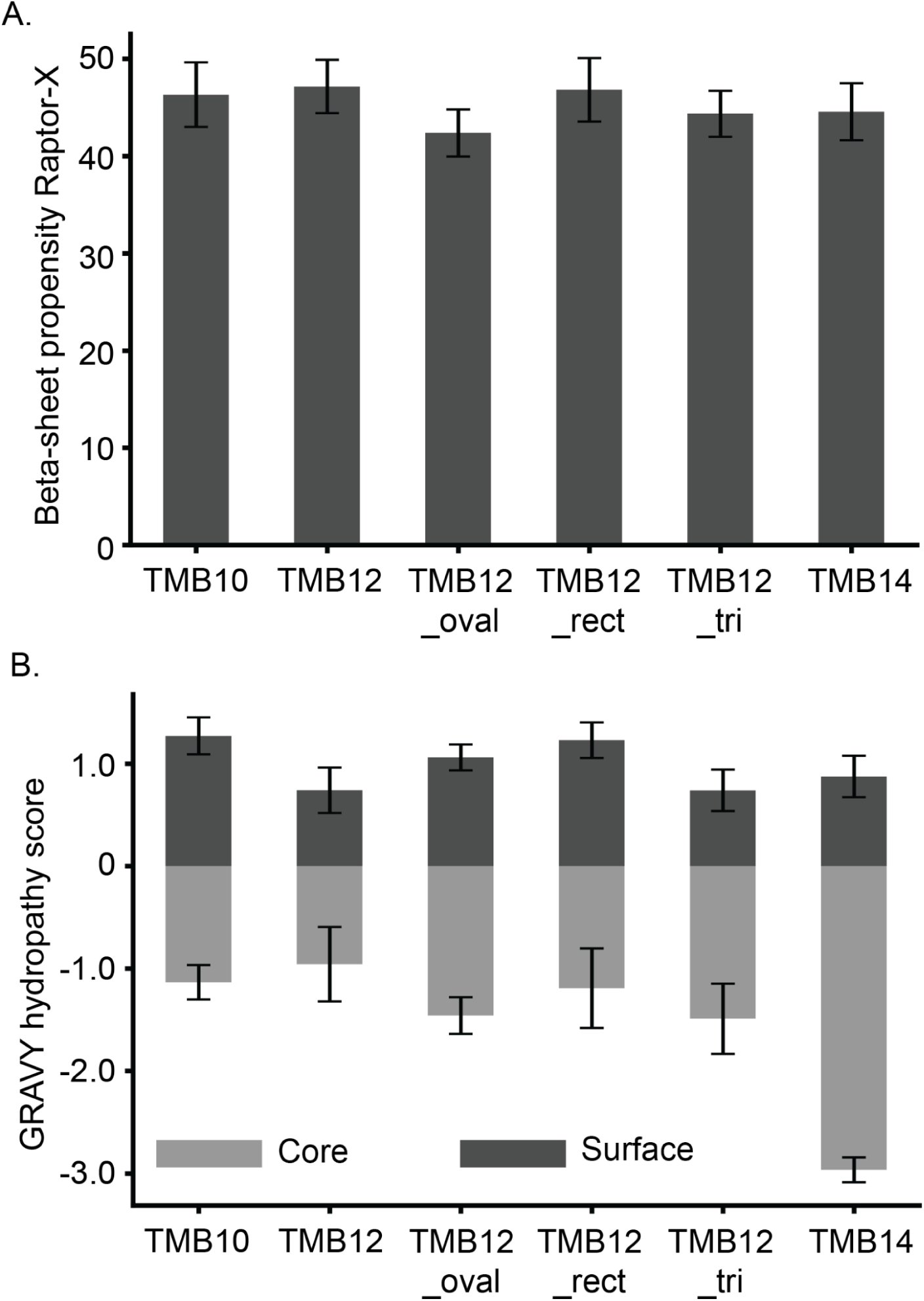
A. Predictions of beta-sheet propensities using RaptorX for different designs. B. GRAVY hydropathy values for the different types of designs and their differences between the pore lining core and surface exposed residues.

**Figure S5:**
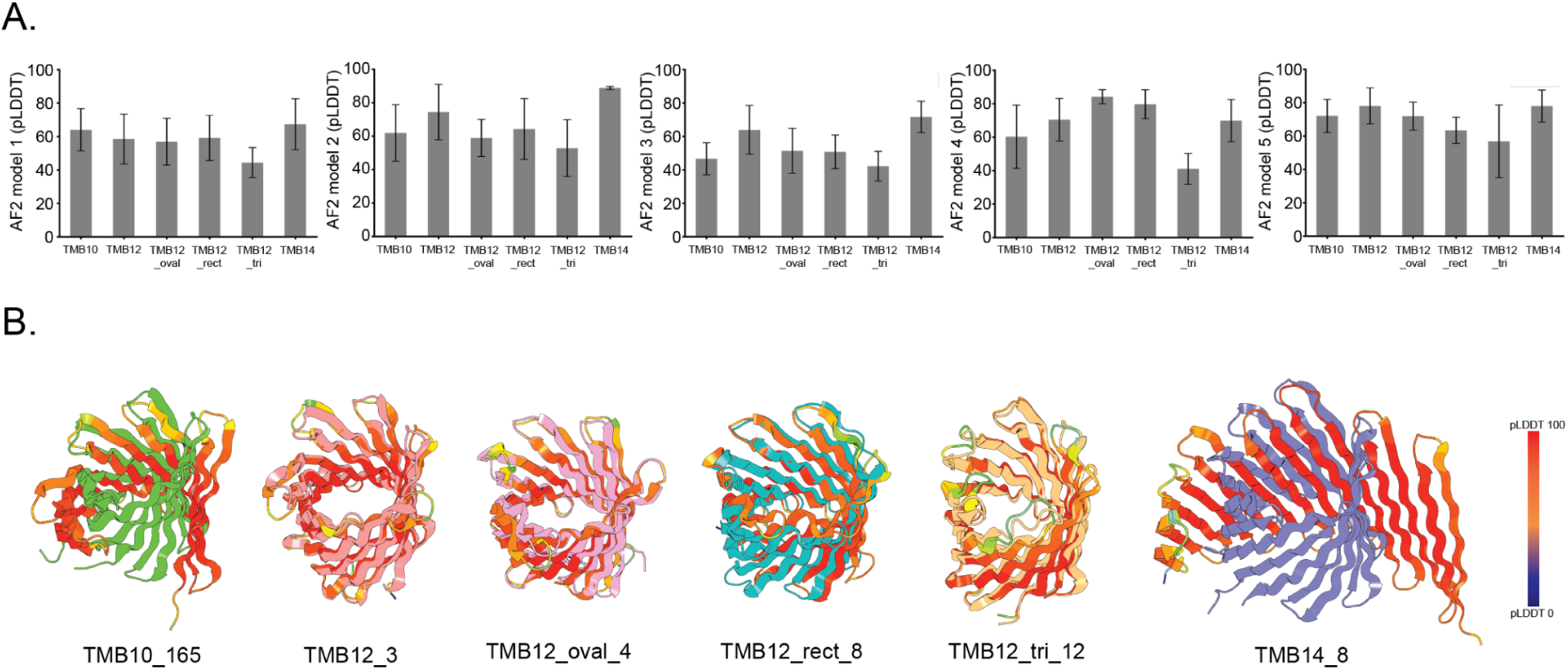
A. Mean pLDDT values for the different types of designs predicted from single sequence without MSA using Alphafold2 and the 5 models. B. Best predicted structures from Alphafold2 for a given type of design (shown in color). The Alphafold2 structures are colored by per residue pLDDT values.

**Figure S6:**
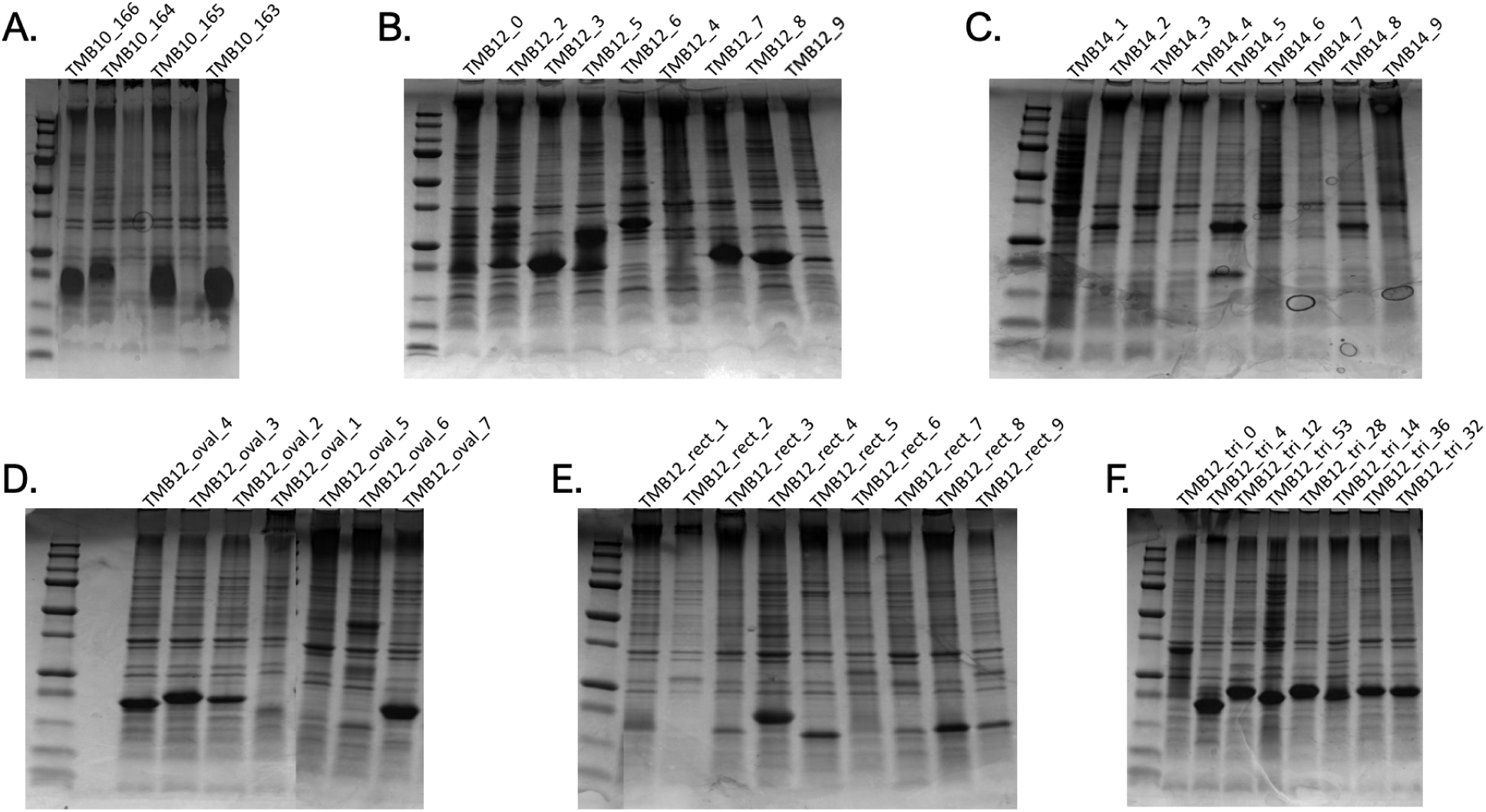
Coomassie stained SDS-PAGE gels showing expression bands for the different designs from insoluble fractions of corresponding lysed cell pellets.

**Figure S7:**
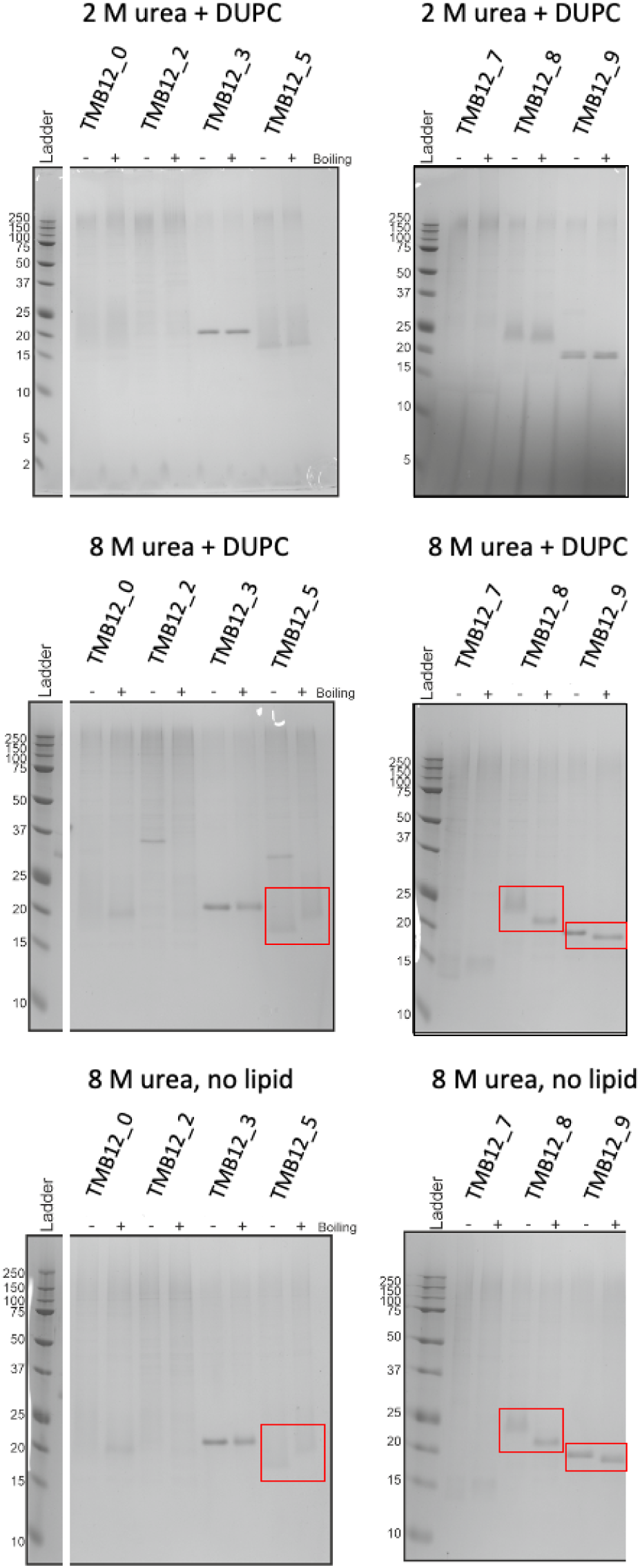
De novo designed TMBs do not exhibit the heat-modifiable behavior on cold SDS-PAGE gel characteristic of folded natural TMBs. No shift in band positions were observed after boiling the samples refolded in DUPC large unilamellar vesicles (LUVs). Band shifts after boiling (red rectangles) were observed only in conditions where the native β-barrel fold can not form (8 M urea with or without lipids).

**Figure S8:**
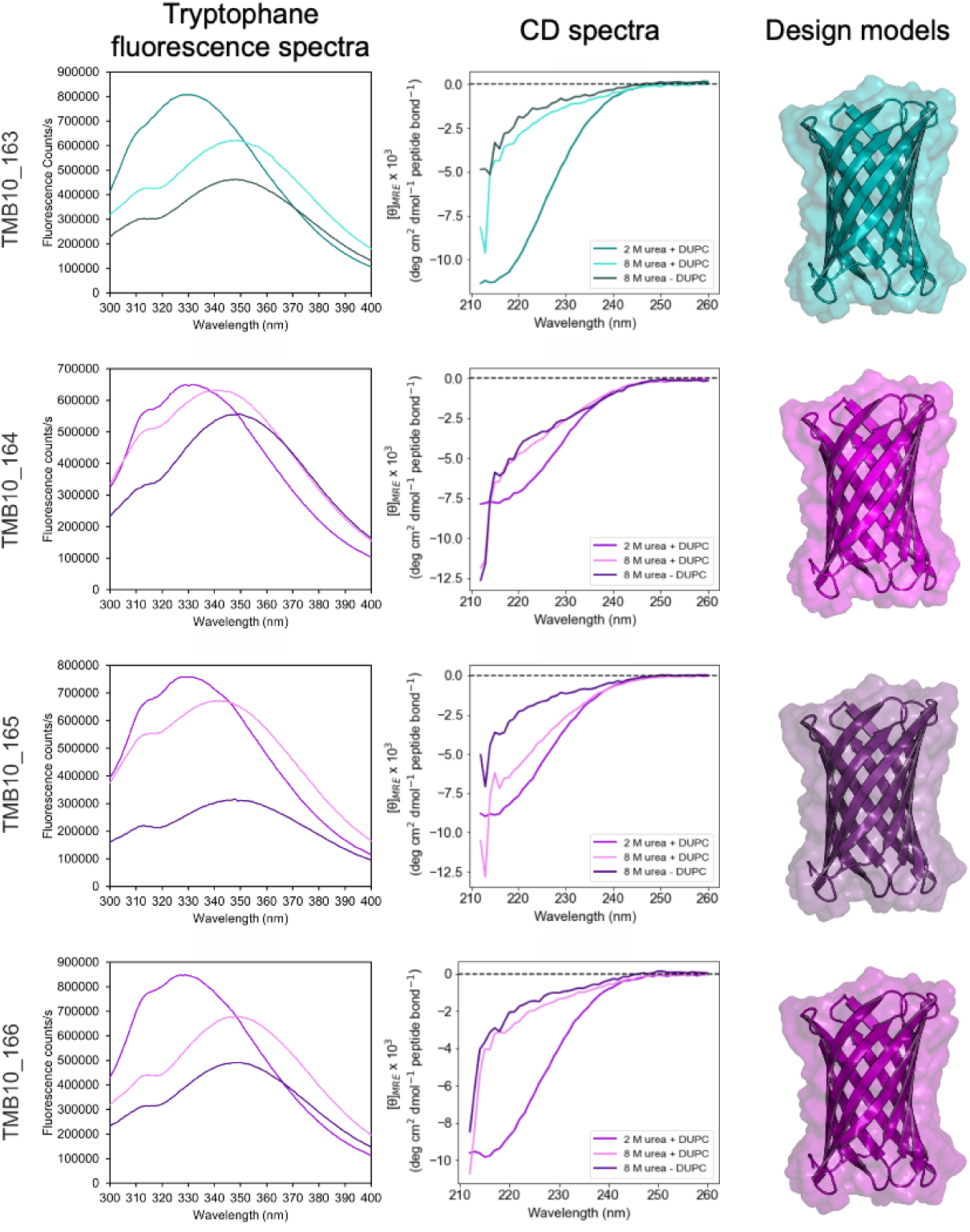
Biophysical characterisation of TMB10 designs (right: design models) for folding in DUPC LUVs. Tryptophan fluorescence spectra (left) and far-UV CD spectra (center) are shown for re-folded proteins in the presence of LUVs and 2M urea (allows TMB folding while reducing aggregation in water), in 8 M urea in the presence of LUVs and in 8 M urea in the absence of lipids. TMB10_163 was selected for further characterization (teal), as it demonstrated a clear fluorescence λ_max_ shift and change in β-sheet structure content between 2 M urea, 8 M urea, and no lipid conditions.

**Figure S9:**
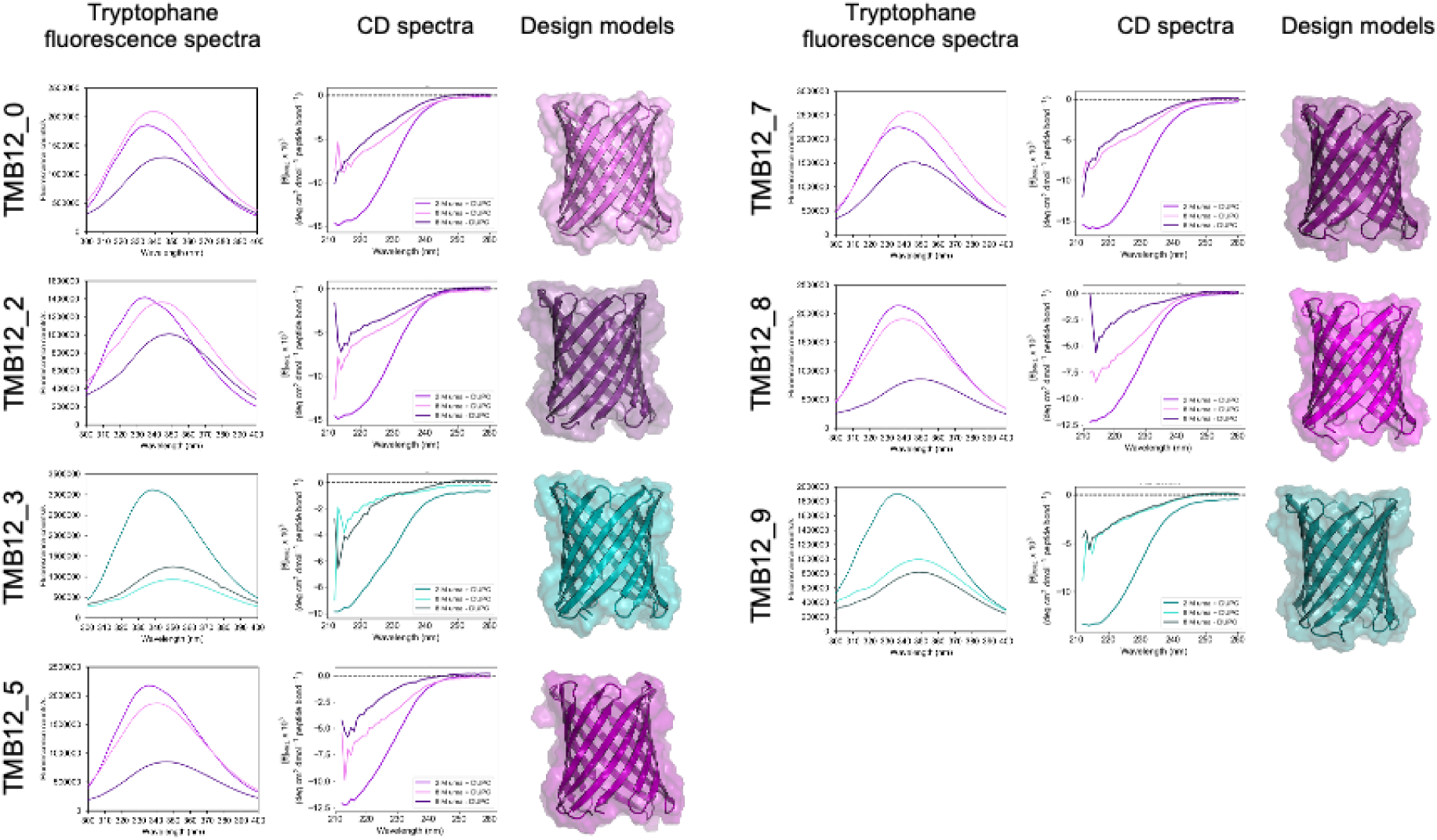
Biophysical characterisation of TMB12 designs with a square shape (right: design models) for folding in DUPC LUVs. Tryptophan fluorescence spectra (left) and far-UV CD spectra (center) are shown for re-folded proteins in the presence of LUVs and 2M urea (allows TMB folding while reducing aggregation in water), in 8 M urea in the presence of LUVs and in 8 M urea in the absence of lipids. TMB12_3 was selected for further characterization (teal), as it demonstrated a clear fluorescence λ_max_ shift and change in β-sheet structure content between 2 M urea, 8 M urea, and no lipid conditions. Similar spectra are observed for the TMB12_9 design, suggesting that the design is also folding into a TMB structure.

**Figure S10:**
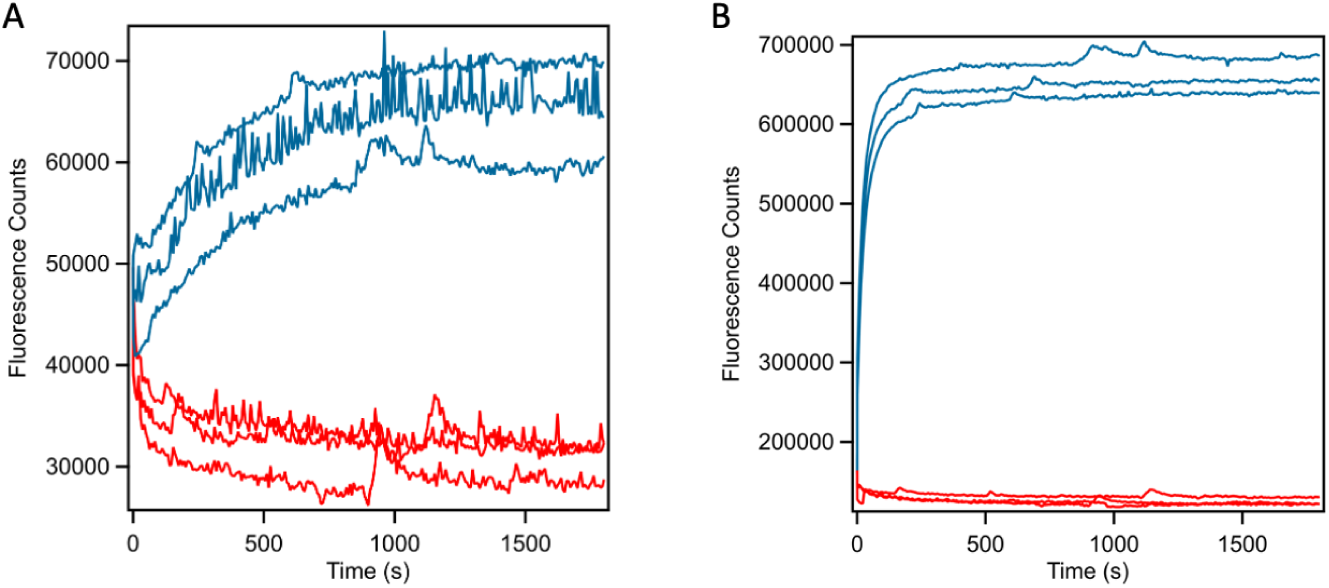
Folding kinetics of designs TMB10_163 (A) and TMB12_3 (B) in DUPC (blue lines) and DMPC (red lines) LUVs at 30°C and monitored by intrinsic tryptophan fluorescence. The difference in folding rates associated with the length of the lipid chain is consistent with intramembrane folding.

**Figure S11:**
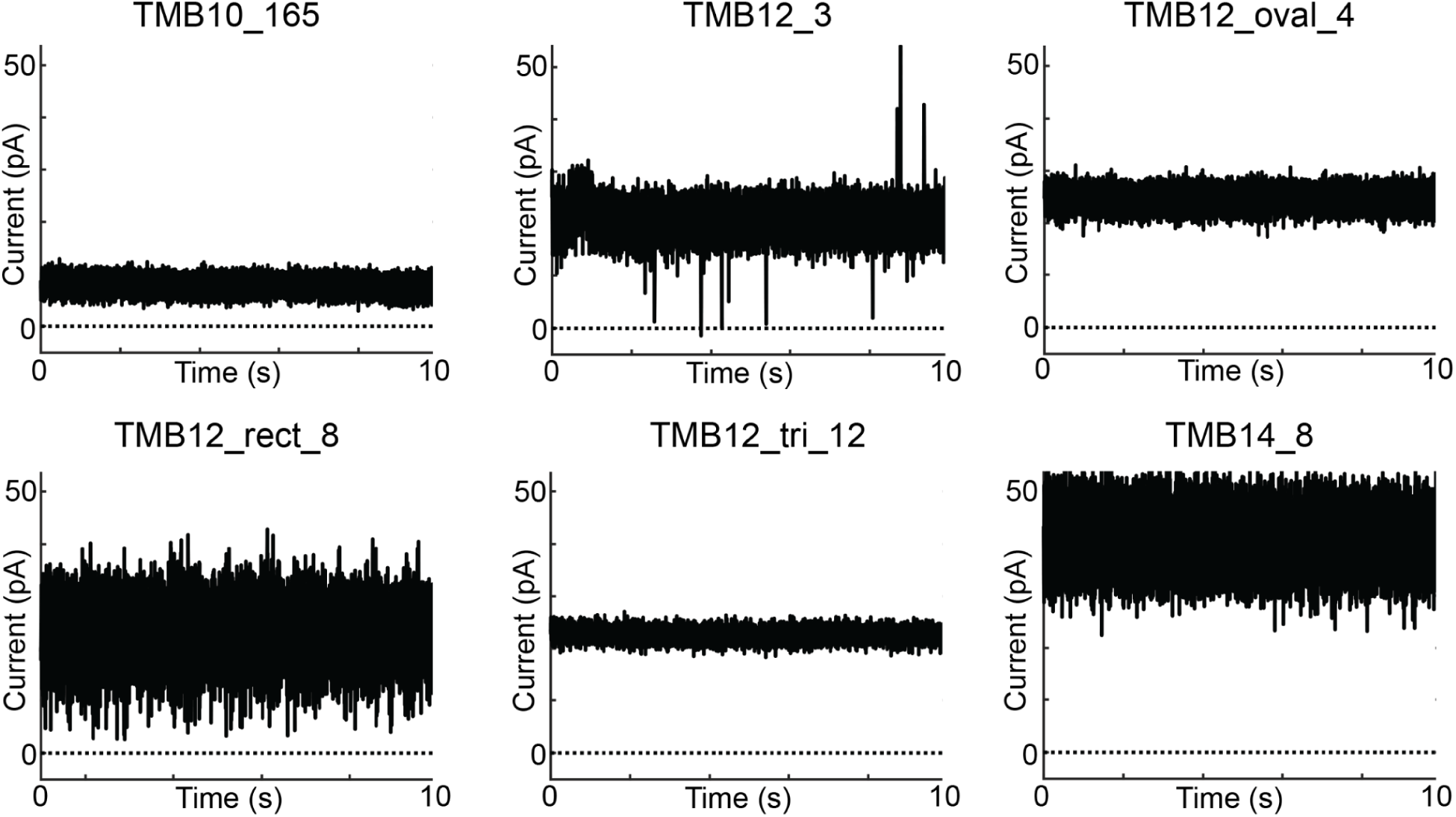
Raw unfiltered current traces of one example for each different type of pores recorded at 5kHz sampling rate. Applied voltage is 100 mV and the cis and trans buffer for all conditions is 500mM NaCl. Different noise levels are a result of the different bilayer capacitances at the time of recording and noise from adjacent cavities in the MECA recording chip from Nanion. 10s reads show characteristics of stable non-gating pores in the membrane.

**Figure S12:**
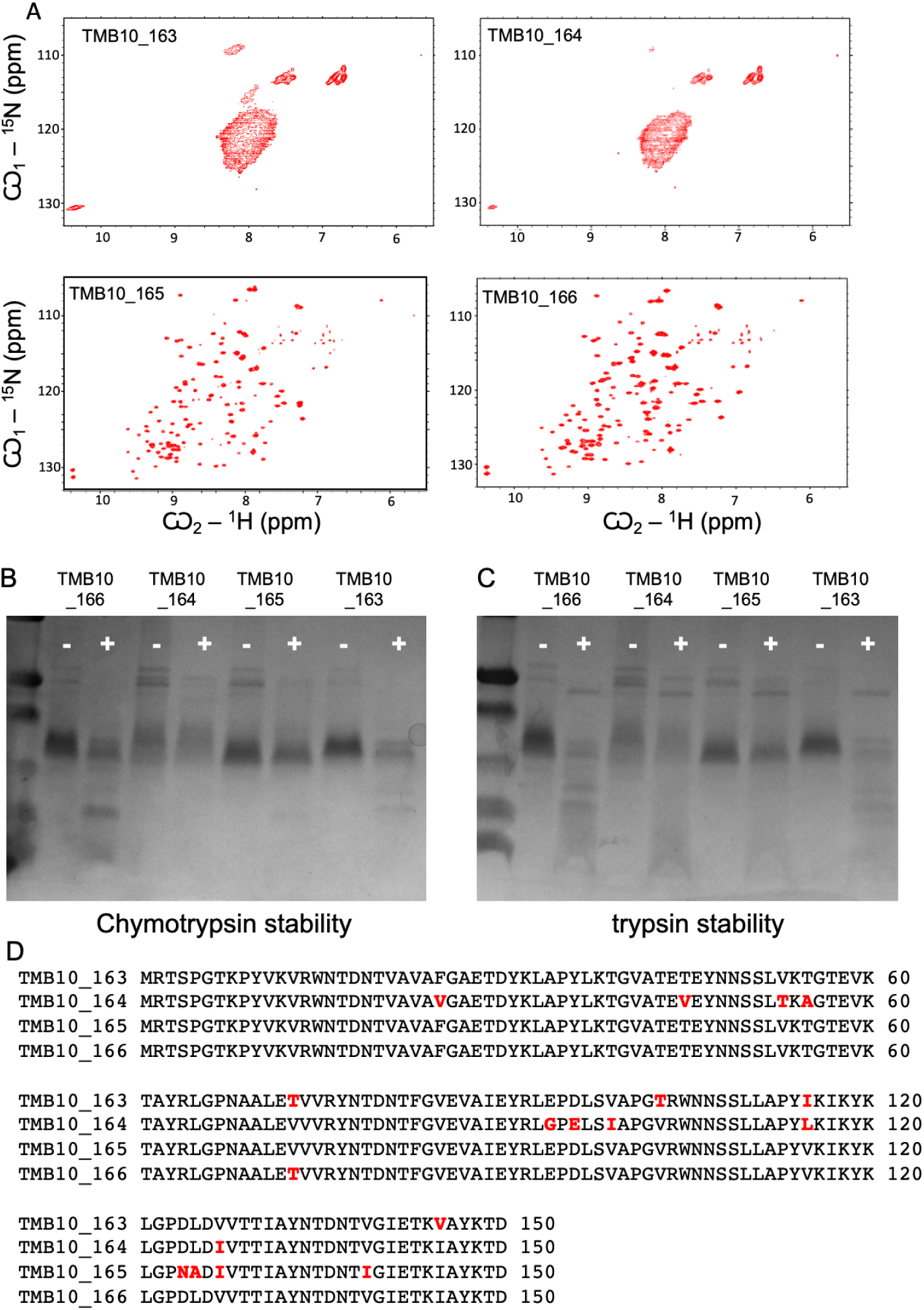
Relative stability of TMB10 designs (163-166). (A) Designs TMB10_165 and TMB10_166 feature well dispersed NMR ^1^H-^15^N HSQC spectra. (B-C) Trypsin and chymotrypsin challenge reveals differences in stability between designs, with TMB10_165 showing the highest stability to both proteases. (D) The designs differ only by 2-9 residues on the lipid-exposed surface, introduced using Rosetta.

**Figure S13:**
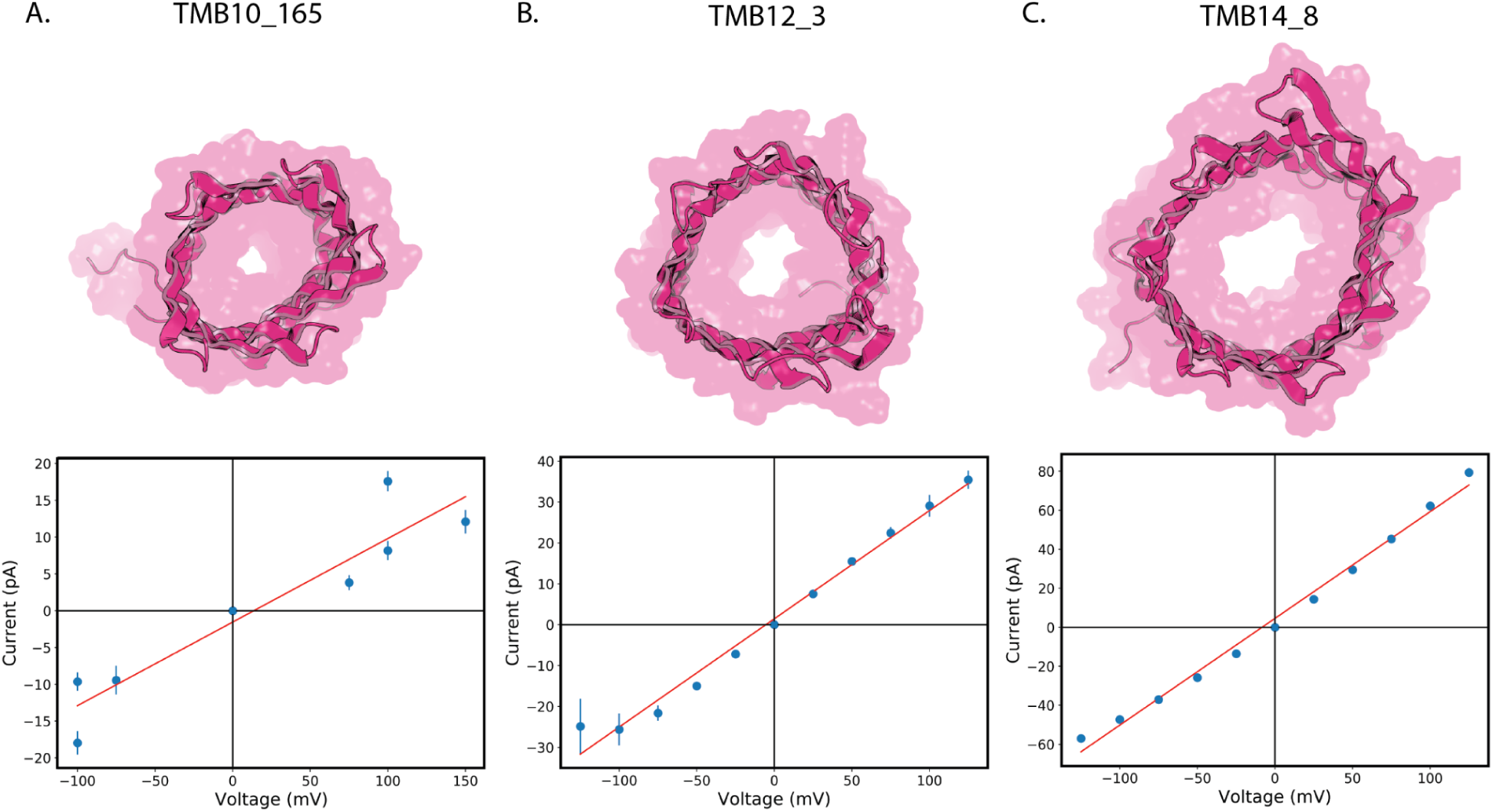
Current vs Voltage plots for three designs pertaining to TMB10_165, TMB12_3 and TMB14_8 pores. All measurements were carried out in 500mM NaCl solution (symmetric across bilayer).

**Figure S14:**
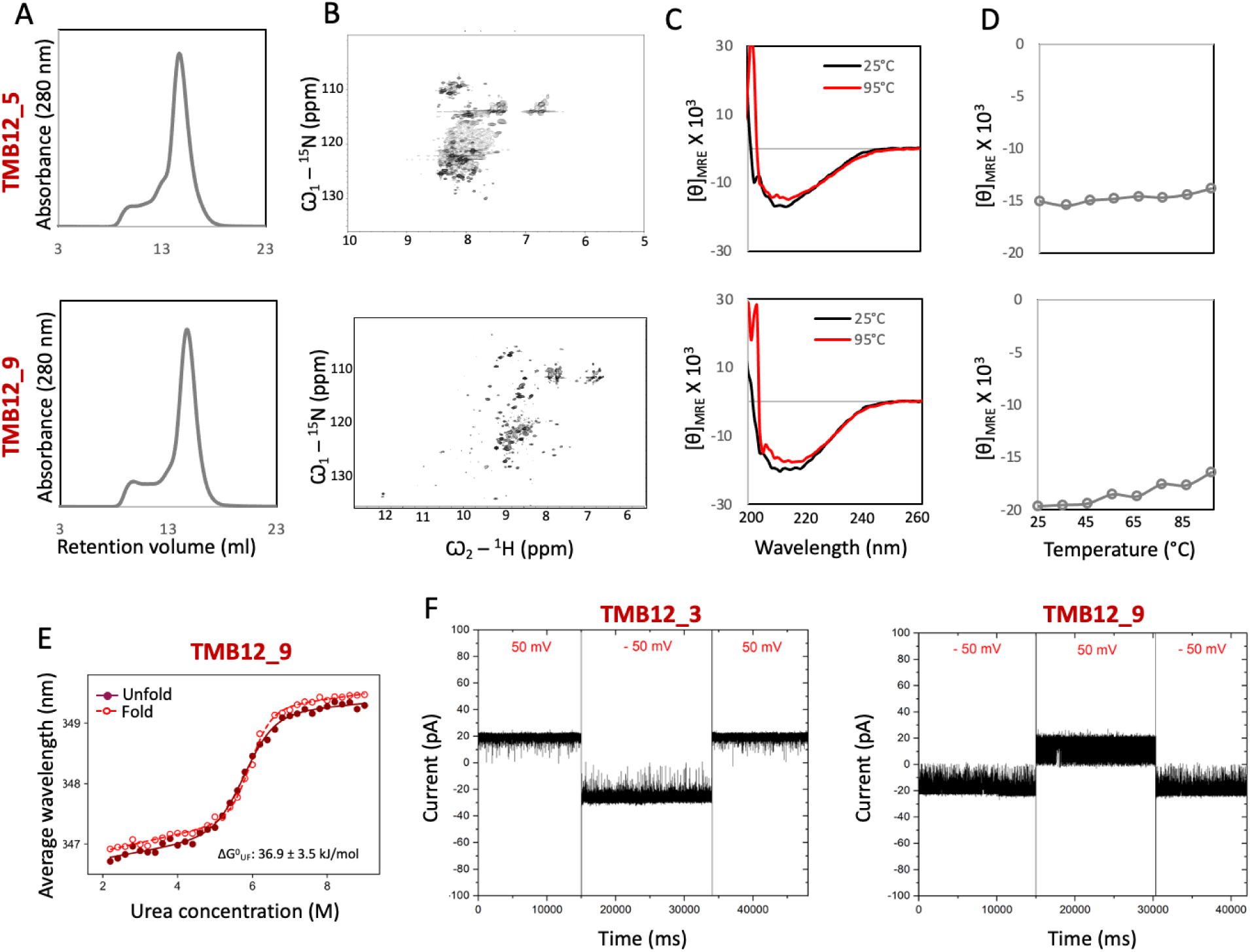
Designs TMB12_5 and TMB12_9 both feature SEC elution profiles consistent with predominantly monomeric TMB12 designs (A). However, only TMB12_9 has a dispersed NMR ^1^H-^15^N HSQC spectrum in DPC micelles indicative of a folded TMB (B), despite yielding far-UV CD spectra characteristic of β-sheet proteins in DPC micelles (C) that remain stable up to 95°C (D). TMB12_9 cooperatively and reversibly folds/unfolds in DUPC LUVs with a similar Cm^F^ to TMB12_3 (5.7 ± 0.3 M) but with a less sharp transition and hence lower unfolding free energy (E). Stable nanopore activity was observed for designs TMB12_3 (stable signal) and TMB12_9 (gated signal suggesting lower stability) (F). No nanopore activity could be observed for TMB12_5, despite several attempts to insert it into DphPC membranes.

**Figure S15:**
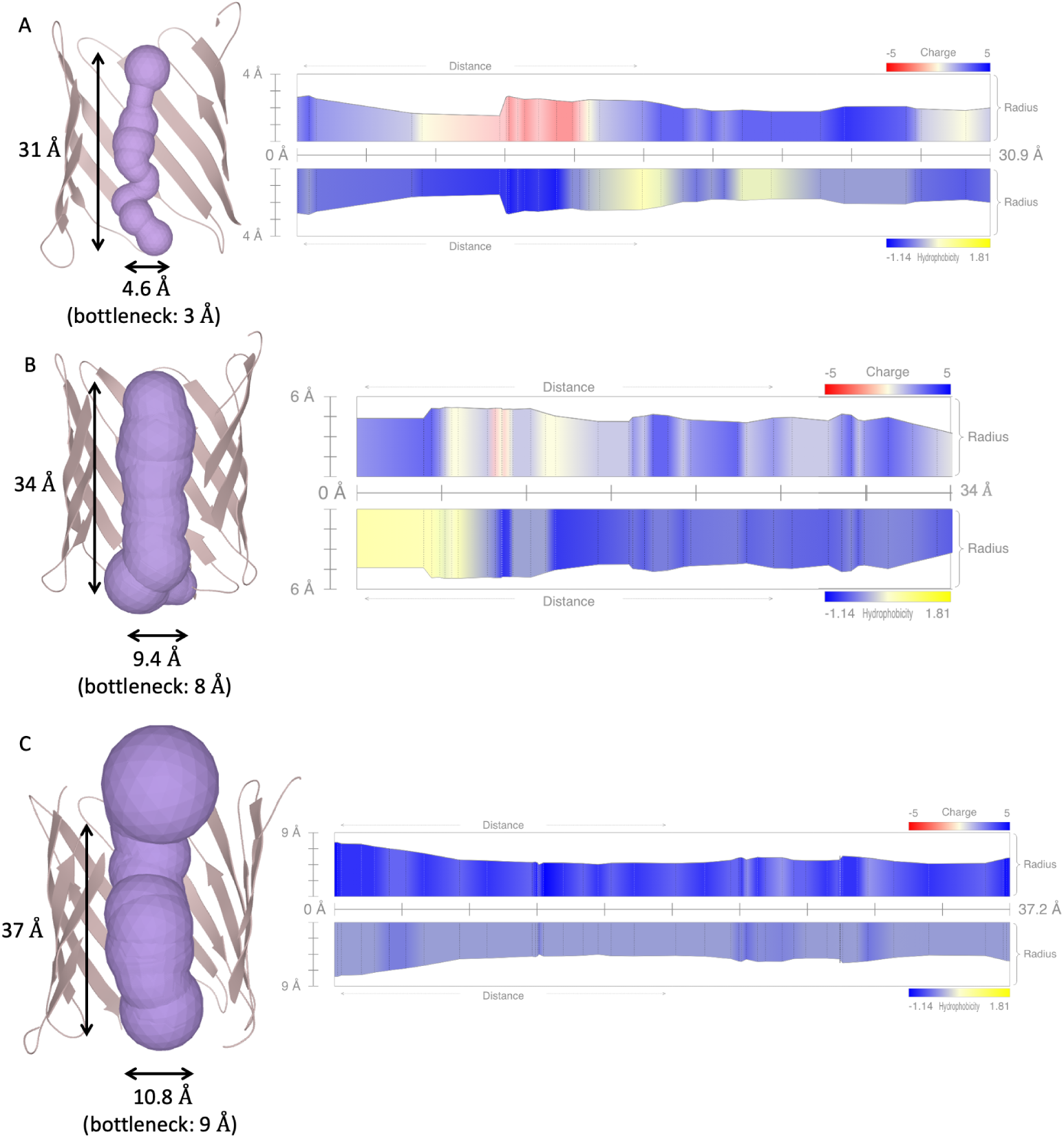
MOLE 2.5 pore size calculations (left), charge and hydrophobicity profiles (right) for designs TMB10_165 (A), TMB12_3 (B) and TMB14 (C).

**Figure S16:**
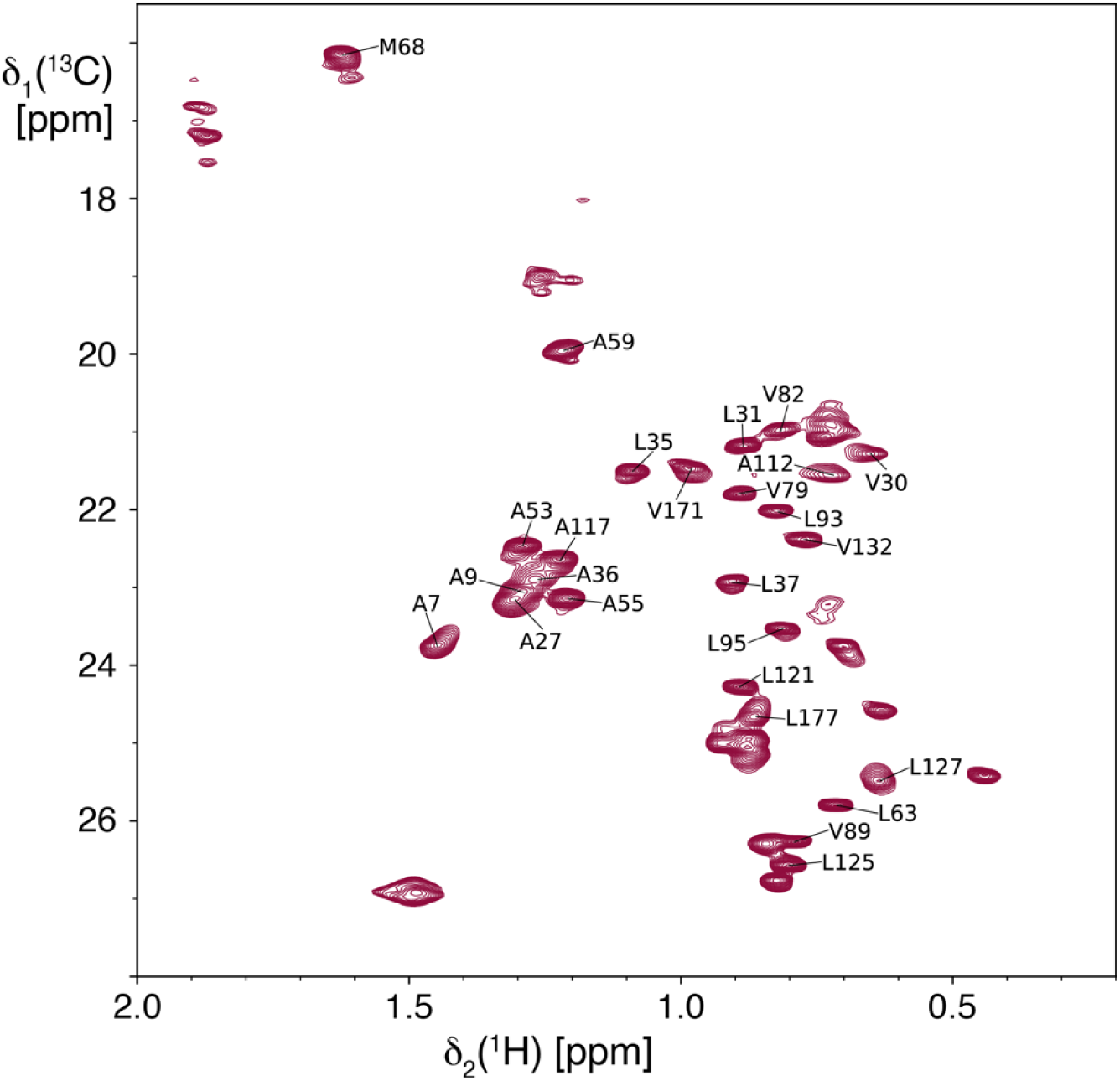
2D [^13^C,^1^H]-HMQC of AMLV-^1^H^13^C-methyl-labeled TMB12_3 in LDAO. Sequence-specific resonance assignments are indicated.

**Figure S17:**
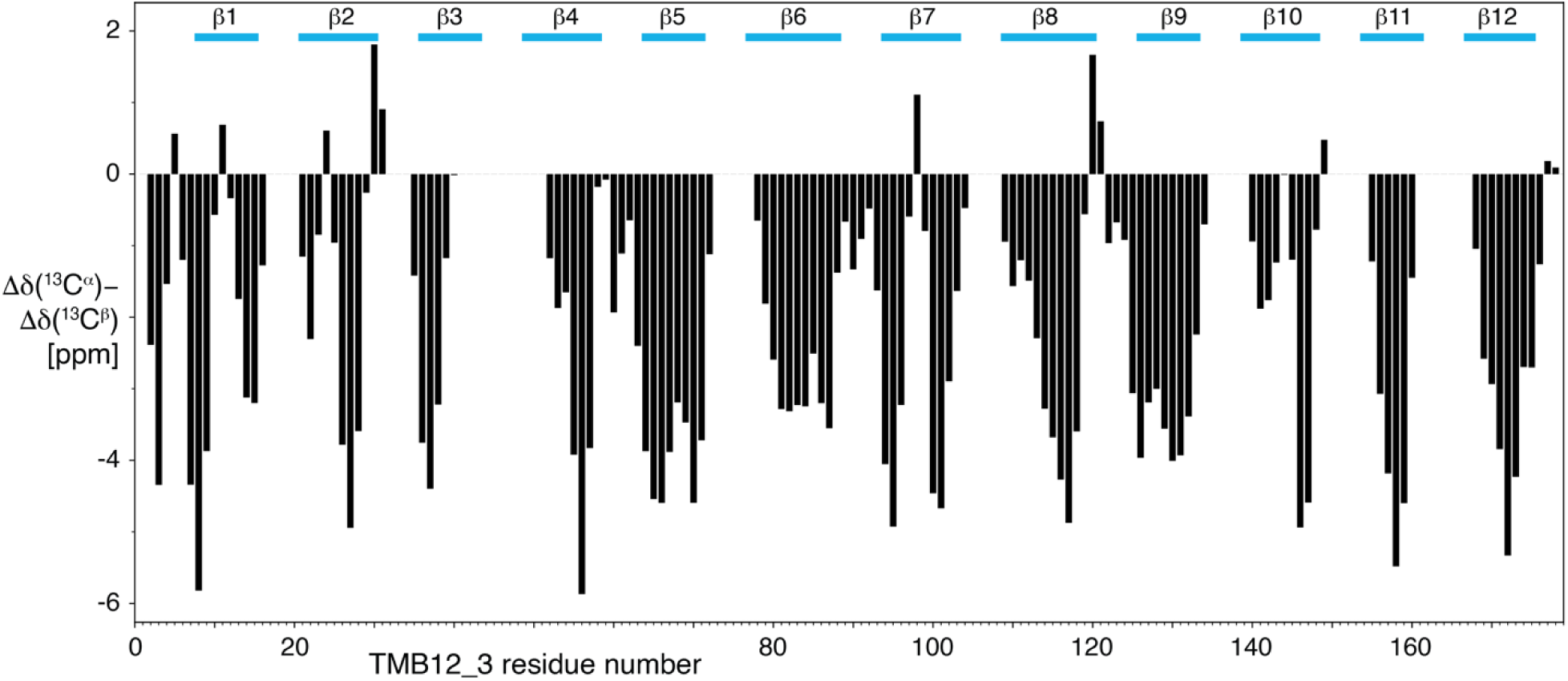
Secondary chemical shifts of TMB12_3 from sequence-specific resonance assignments. Consecutive stretches of large negative values indicate the presence of β-strand secondary structure. The positions of the 12 β-strands are indicated by blue lines.

**Figure S18:**
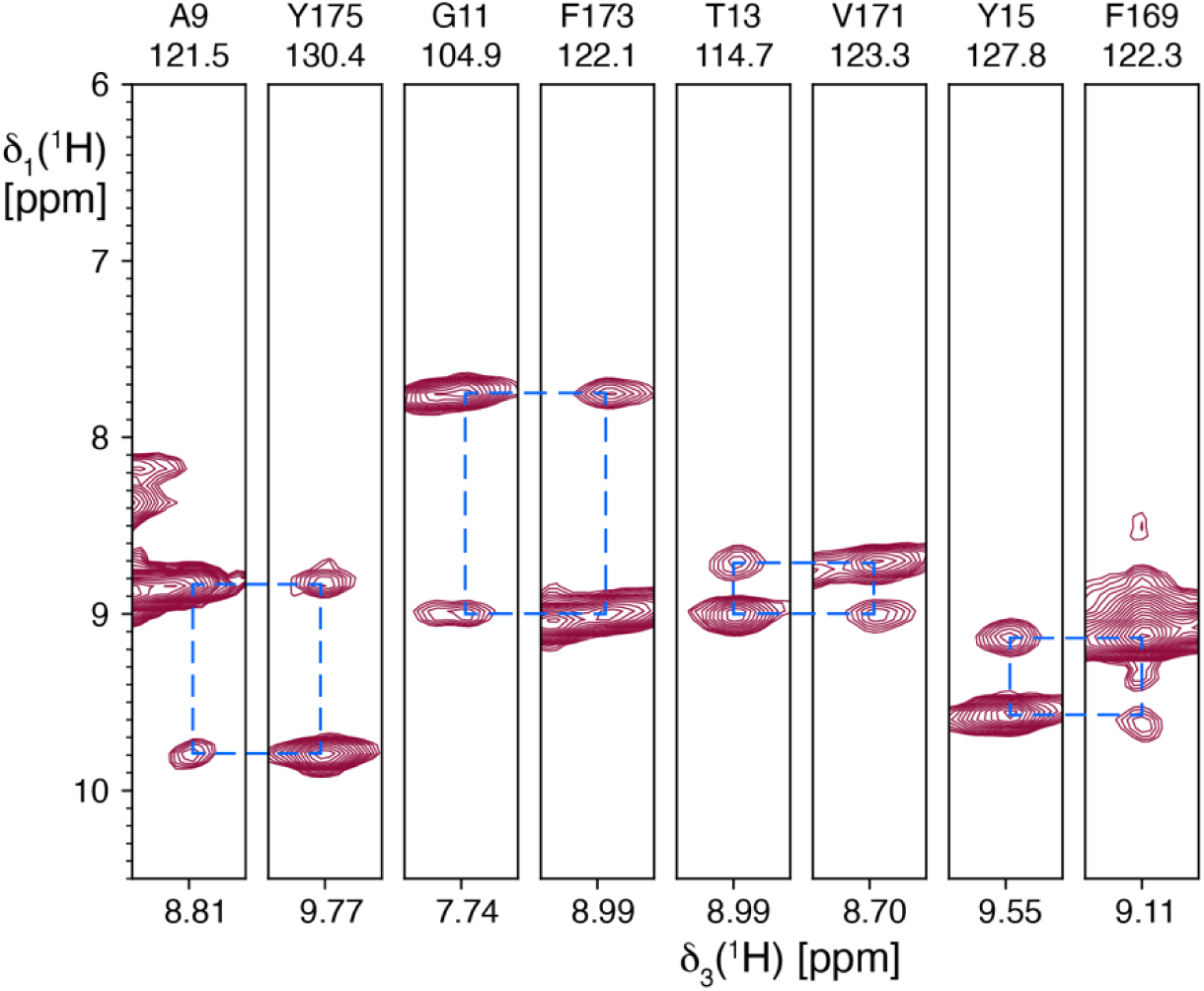
Strips from the 3D [^1^H,^1^H]-NOESY-^15^N-TROSY experiment of TM12_3 in LDAO micelles. Strips were taken for the residue pairs involved in the antiparallel β1–β12 pairing. The NOE cross peaks are connected to the diagonal peaks by blue dashed lines.

**Figure S19:**
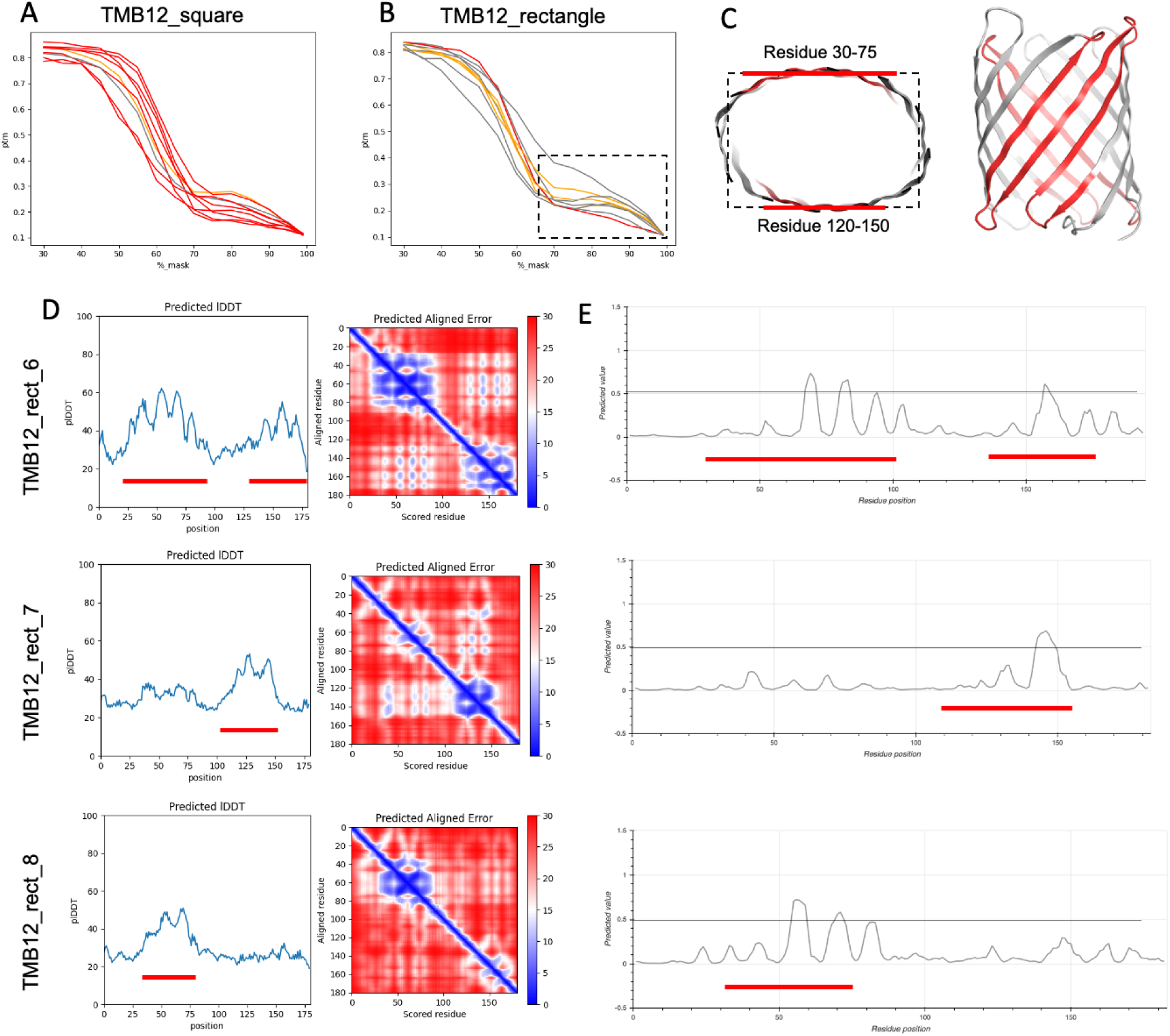
ESM noise ramp (*in silico* melting (*21*) simulations) plots for (A) square-shaped TMB12 designs and (B) rectangle-shaped designs (curves colored by protein expression level, as defined in Figure S6. red=strongly expressed; orange=weakly expressed; gray=no expression). The rectangle-shaped designs feature more residual secondary structure at the end of simulations (highlighted by a dashed rectangle). Closer analysis of the ESM simulations show that the regions of melting-resistant secondary structure correspond to the long sides of the designed rectangular TMB12 structures (C). These regions (highlighted with red lines) have higher plDDT scores (D, left) and lower pAE scores (D, right) after 85 % of the sequence is masked to the ESMFold language module. They co-localize with early-folding regions predicted with EFoldmine (*22*) (early-folding score surpassing the 0.5 threshold, highlighted with red lines) (E).

**Figure S20:**
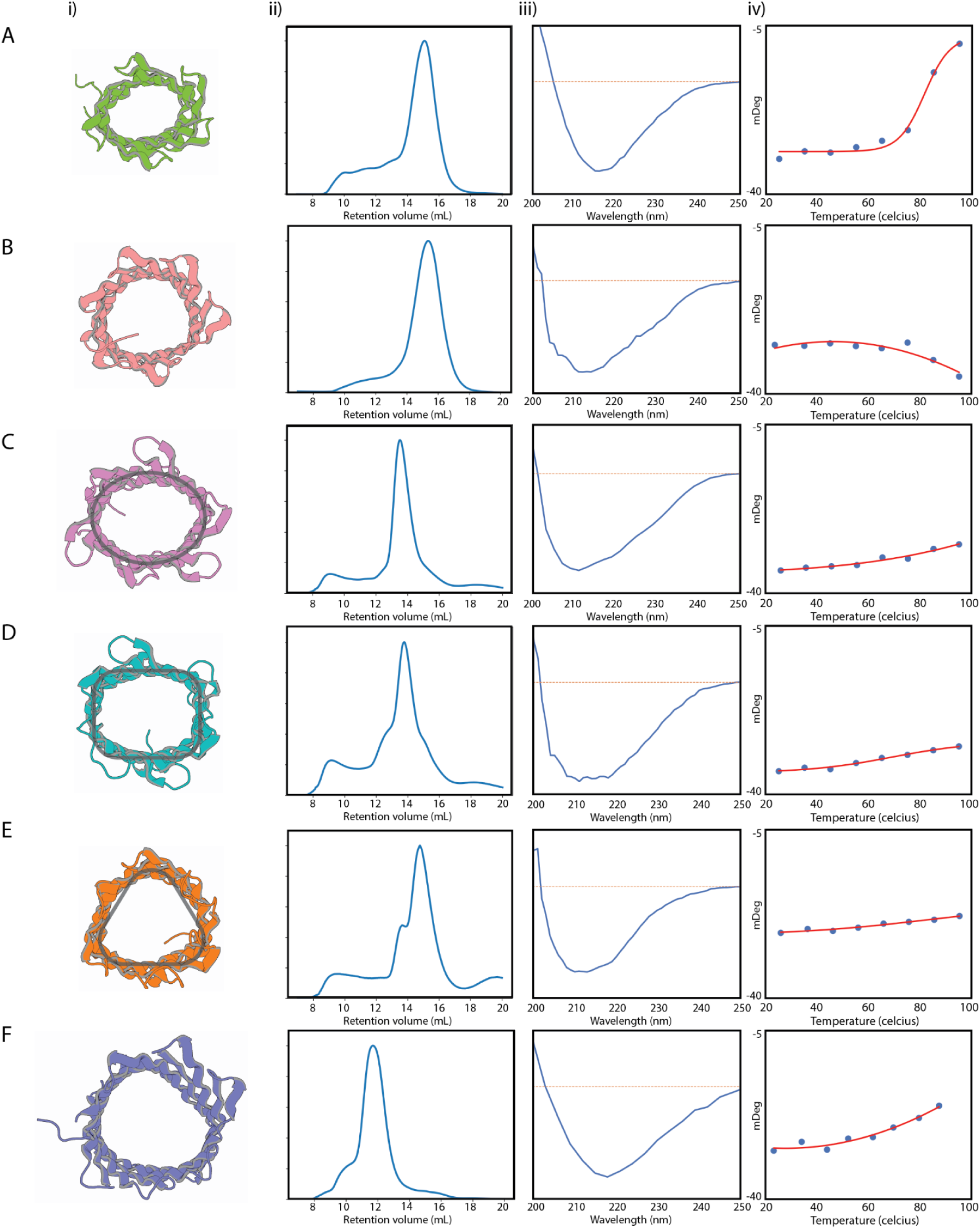
Characterisation of designed TMB pores. A. TMB10_165, B. TMB12_3, C. TMB12_oval_4, D. TMB12_rect_8, E. TMB12_tri_12, F. TMB14_8. i) Respective cartoons indicating top view of the designs. ii) Size Exclusion Chromatography plots for all designs carried out in a buffer containing 0.1% w/v DPC detergent. iii) Corresponding CD plots and iv) temperature ramp plots from 25°C to 95°C.

**Figure S21:**
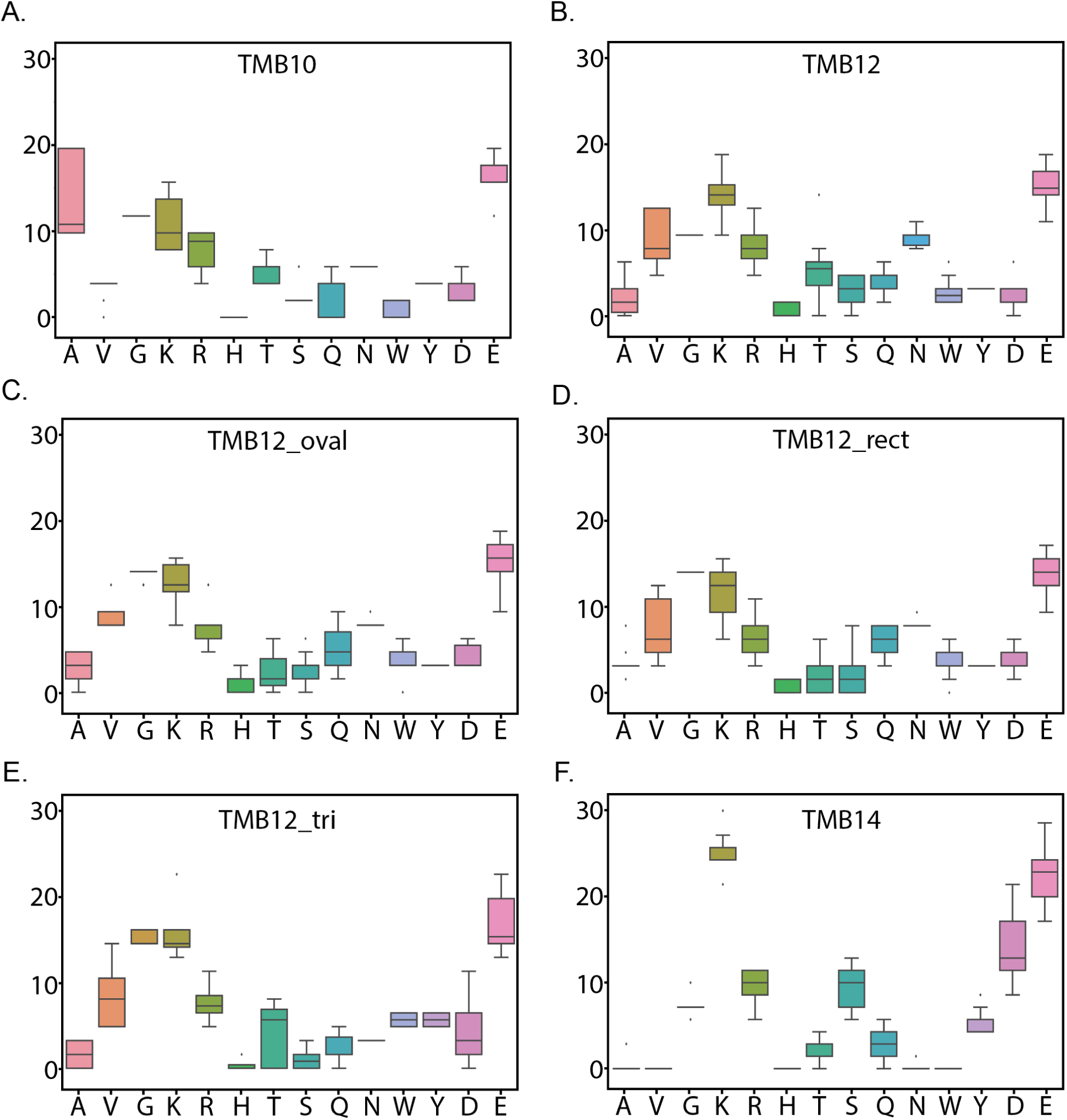
Pore lining amino acid compositions of the different types of designs. Y-axis indicates the percent fraction of the total number of a specific amino-acid within all pore-lining residues.

